# Mutational spectra reveal influenza virus transmission routes and adaptation

**DOI:** 10.1101/2025.11.26.690773

**Authors:** Christopher Ruis, Artur Duque Rossi, Theo Sanderson, Marta Matuszewska, Maria Ntemourtsidou, Vítor Mendes, Pedro Henrique Monteiro Torres, Julian Parkhill, Thomas P. Peacock, R. Andres Floto

**Author notes:** Correspondence to: Christopher Ruis or Andres Floto.

## Abstract

Influenza A virus remains a major cause of morbidity and mortality in humans and animals with pandemic potential. While control of outbreaks and spillover events remains a key health priority, the transmission routes and adaptations causing these episodes remain unclear. To understand these processes, we compared mutational spectra across the complete diversity of influenza A virus. We find that niche-specific mutagens cause large convergent shifts in mutational spectrum between gastrointestinal and respiratory lineages, allowing inference of transmission route and site of infection, while additional mutational patterns are host-specific, permitting detection of species responsible for outbreaks. By identifying points of spectrum change in phylogenetic trees, we discover novel adaptive mutations enabling sustained respiratory transmission. We conclude that mutational spectra can enable early detection of lineages with increased potential for spillover and onward transmission, and should be considered as a component of genomic surveillance strategies.

## Main Text

Influenza A virus (IAV) causes roughly one billion cases and up to 650,000 deaths annually in humans (*1*), is responsible for tens of millions of deaths and billions of dollars of associated economic costs across commercial poultry and swine populations (*2–6*), drives outbreaks within companion and farm animals (*7*, *8*), and is emerging as a key driver of morbidity and mortality in a broad diversity of wild mammals and waterbirds (*9*). Furthermore, IAV has caused at least six human pandemics since 1889, associated with high infection rates and vast excess deaths (*10*), and remains a major risk for future pandemics (*9*, *11*). All influenza lineages within mammals and poultry ultimately originate from a highly diverse viral reservoir within wild waterbirds of the orders Anseriformes (including ducks, geese and swans) and Charadriiformes (including gulls and shorebirds) (*12–14*). We currently have an incomplete understanding of IAV transmission amongst wild waterbirds, how IAV jumps between species, and the key steps involved in virus adaptation to undergo sustained transmission within a new host.

We have recently shown that mutational spectra (the contextual patterns of nucleotide substitution acquired during evolution (*15–17*)) can provide new insights into microbial ecology. We have found that microbes that transmit through different routes and particular hosts are exposed to distinct sets of mutagens which leave specific mutational signatures within microbial genomes (*18–23*). Thus, by calculating and detecting mutational signatures, it is possible to identify dominant transmission routes (*18*, *19*), infer host species driving outbreaks (*22–25*), reveal impacts of antiviral drug therapy (*21*), and rapidly predict virulence of emerging viral lineages (*20*). We hypothesised that large-scale comparison of mutational spectra across IAV lineages from different hosts would identify host-specific mutational signatures that could be applied to infer predominant IAV transmission routes, pinpoint inter-species jumps, and thereby identify mutations associated with virus adaptation. We therefore examined mutational patterns across the entire diversity of IAV to uncover mutational signatures associated with distinct transmission routes and host species.

### Specific patterns of mutation occur during IAV evolution

To examine host-specific mutational patterns, we first calculated the context-specific mutational spectra of 61 IAV lineages (using *MutTui* (*26*)), covering the entire known diversity of haemagglutinin (HA) and neuraminidase (NA) segments (***Figures 1A-B****, **S1A**, **Table S1**, **Supplementary text***). Of these lineages, 21 infect a variety of wild waterbirds, three are associated with gulls (*Laridae*) (*27*), 16 exhibit sustained circulation in specific mammal species, 11 show sustained circulation in gallinaceous poultry, four belong to the High Pathogenic Avian Influenza (HPAI) Goose-Guangdong (GsGd) H5 lineage (*9*, *11*), and six circulate within domestic Anseriformes (***Figure 1A****, **Supplementary text***). In total, we analysed over 430,000 mutations acquired during the evolution of more than 75,000 IAV sequences (***Figure 1C***).

**Figure 1.**
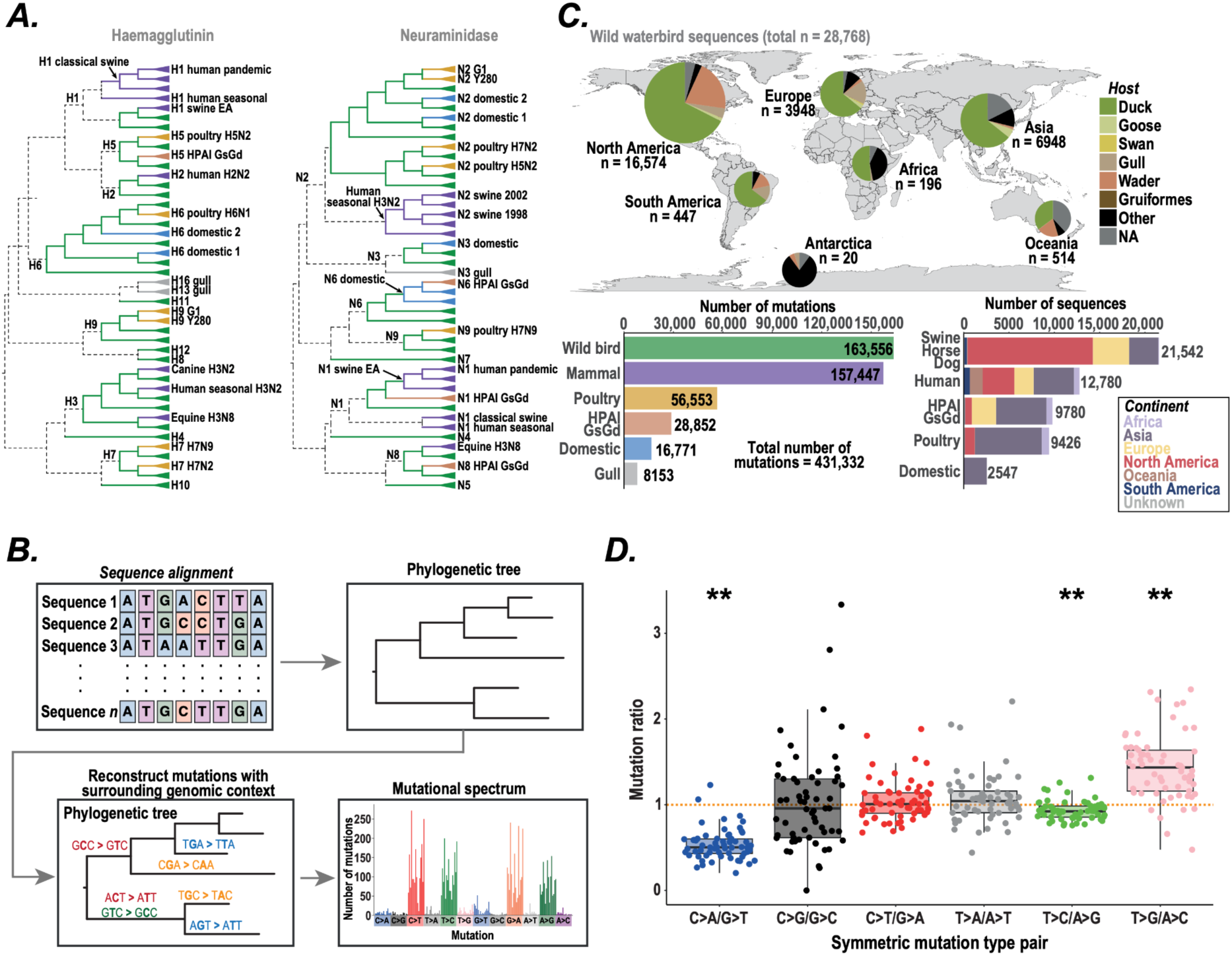
The mutational patterns of IAV. (**A**) Cladograms showing evolutionary relationships between IAV lineages for which mutational spectra were calculated and compared. Phylogenetic branches are coloured by host group and triangles on tip branches represent downstream diversity. Subtype labels on internal phylogenetic branches relate to all downstream branches until the next label. Dashed branches were not included within any calculated spectra. (**B**) To calculate a lineage mutational spectrum, we reconstruct a phylogenetic tree of aligned sequences from the lineage. Mutations are reconstructed onto the tree and the surrounding nucleotide context of each mutation identified within the reconstructed sequence at the start of the respective branch. The number of occurrences of each contextual mutation is counted to calculate the mutational spectrum. The mutational spectrum shown is that of H1 in wild waterbirds. (**C**) The map shows the geographical distribution of wild waterbird sequences coloured by host group. The lower right panel shows the number of sequences included in non-wild waterbird datasets coloured by continent, while the lower left panel shows the number of sampled mutations within each host group. (**D**) We compared the frequency of symmetric mutation type pairs across rescaled mutational spectra from all lineages. Mutation pairs where the ratio is significantly different from one, showing elevation of one mutation type over the other, are indicated (** shows Benjamini-Hochberg corrected *p* < 0.01).

We found that IAV mutational spectra are consistently dominated by transition mutations (median proportion 0.83, range 0.73-0.91, ***Figure S1***) (*28*, *29*), with C>T and G>A mutations occurring at the highest rates once spectra are rescaled by genome composition (ANOVA *p* < 0.05, ***Figure S1***, see ***Methods***). We identified G>T as the most common transversion mutation, consistent with virus exposure to reactive oxygen species (*20*, *30*). We observed that the surrounding nucleotides influence the substitution frequency of each base change (Benjamini-Hochberg-adjusted *p* < 0.05, ***Figure S1***). While some contextual patterns were consistent with selection to avoid the host zinc-finger antiviral protein (ZAP) (*31*), such as elevations in mutations that remove CG dinucleotides and reductions in T>C and A>G transitions introducing CG dinucleotides, we found other context-specific patterns unlinked to ZAP evasion, indicating the action of context-dependent mutational processes on viral genomes (***Figure S1***).

To explore the impact of mutagens, we examined the level of symmetry in base changes, since mutations introduced by the viral polymerase during genome replication are likely to be symmetric (6), while those caused by mutagens (which predominantly target the negative-sense genomic strand) are mostly asymmetric (*20*). We found elevations in G>T over C>A (2-fold), T>G over A>C (1.4-fold), and A>G over T>C (1.1-fold), consistent with a major contribution of mutagens to IAV mutational patterns (***Figure 1D***, ***Figure S2***).

### Transmission route is a major determinant of IAV mutational spectrum

We next sought to identify host-associated mutational signatures. We initially examined spectrum changes across six independent jumps of IAV from wild waterbirds into either mammals or poultry. We found that these host jumps were all associated with convergent, large-scale shifts in mutational types, with significant changes in eight out of the 12 possible base changes in each case (***Figures 2A***, ***S3***). These spectrum shifts were detectable whether we examined spectra that were unscaled, rescaled for genomic composition, or calculated using only synonymous mutations (***Figures 2B***, ***S4***) and were large enough to separate the overall mutational spectra of mammals and poultry away from wild waterbirds (***Figures 2C*, *S5***).

**Figure 2.**
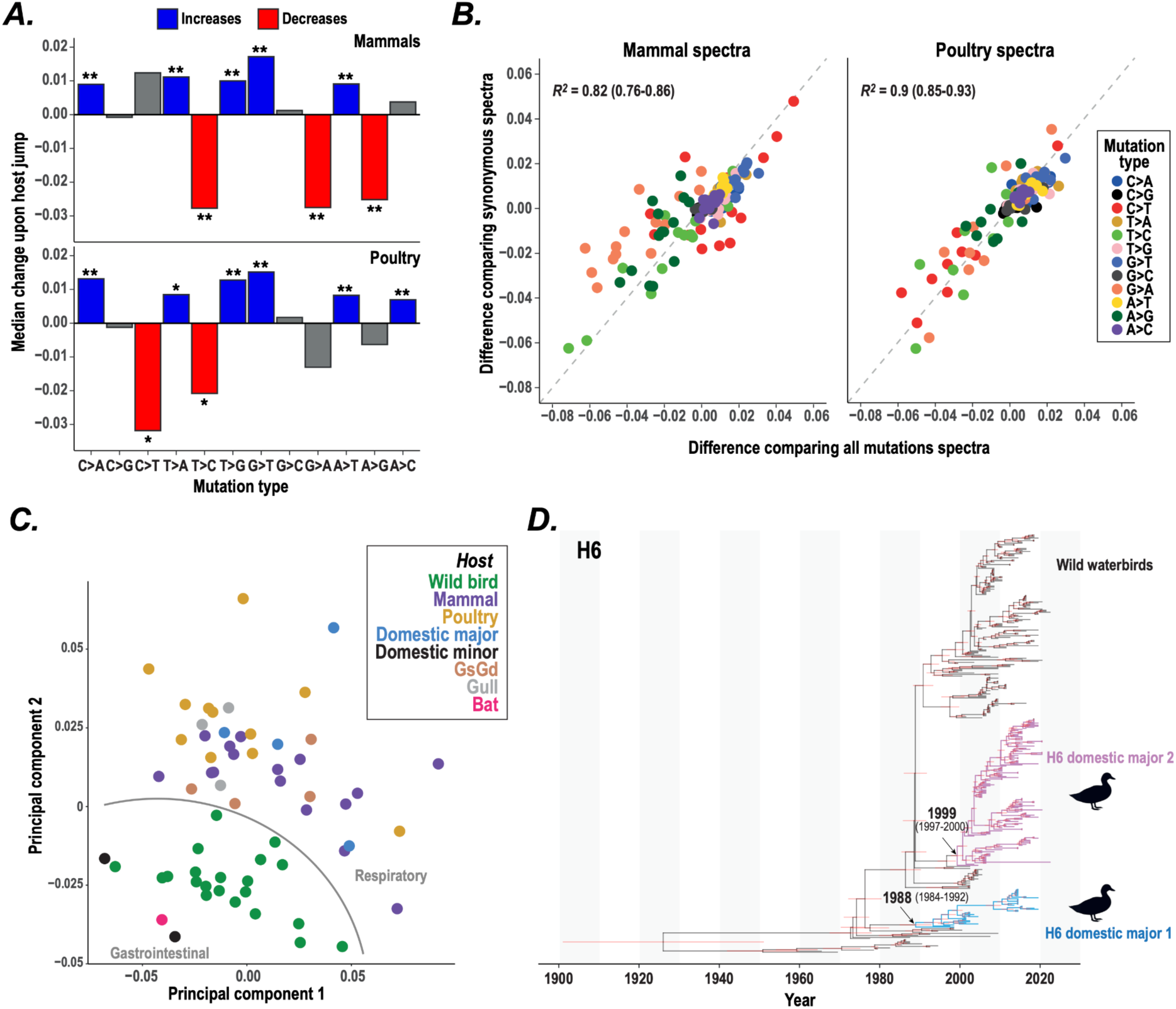
Gastrointestinal and respiratory transmission are associated with distinct mutational spectra. (**A**) We identified mutational spectrum changes associated with host jumps by subtracting the respective ancestral wild waterbird spectrum from each mammal and poultry spectrum. The average change in proportion of each mutation type in unscaled spectra is shown; changes leading to each individual spectrum are shown in ***Figure S3***. * and ** denote Benjamini-Hochberg adjusted randomisation *p* < 0.1 and *p* < 0.05 respectively. (**B**) The proportion change for each mutation type comparing each mammal or poultry spectrum with its respective ancestral wild waterbird spectrum is shown for unscaled spectra and spectra calculated using synonymous mutations only. There is a strong correlation in each case, showing that these patterns are not being driven by natural selection of amino acid substitutions. (**C**) We carried out a principal component analysis on all lineage spectra based on proportions of the twelve mutation types. There is a separation between gastrointestinal and respiratory lineages. (**D**) Temporal maximum clade credibility phylogenetic tree of the H6 subtype, including sequences from the two major H6 domestic lineages (blue and purple branches, respectively) and wild waterbirds (black branches). Most recent common ancestor dates for the major domestic lineages are shown as medians with 95% highest probability density (HPD) in parentheses. Red bars show 95% HPD for node dates.

We wondered whether the spectrum changes across host jumps could be driven by different transmission routes, as we have previously observed for SARS-CoV-2 and bacteria (*19*, *20*), and hypothesised that the spectrum shifts might be caused by changes in dominant transmission mode of IAV: from gastrointestinal transmission, the dominant route in wild waterbirds (*14*), to respiratory transmission, seen in mammals (*6*, *7*, *32*, *33*) and gallinaceous poultry (*34–36*).

To test this possibility, we examined mutational spectra for IAV lineages associated with other host groups. We found that mutational spectra both of gull-adapted lineages (which undergo aerosol transmission and respiratory infections in ferrets (38, 39), and cause predominantly respiratory disease during experimental gull infections (37)) and of HPAI GsGd lineages (which cause respiratory infections in wild waterbirds prior to systemic spread (40–44)) cluster with spectra from mammal and poultry lineages, while the mutational spectrum of bat IAV (which causes gastrointestinal infections (45–48)) clusters with wild waterbirds, supporting the separation of mutational spectra being based primarily on transmission route rather than virus or host phylogeny (***Figures 2C***, ***S6***).

We then analysed IAV lineages within commercial domestic Anseriforme waterbirds (ducks and geese; ***Figure S7***) which, while genetically similar to wild populations, are farmed at high density in poorly ventilated environments and without access to open water (*37–39*); living conditions which are similar to those of commercial gallinaceous poultry (*40*, *41*) and which likely favour efficient respiratory, rather than gastrointestinal, transmission. We found that two major independent H6 lineages continuously circulating for multiple decades within domestic Anseriformes (***Figure 2D***) have mutational spectra that cluster with those from mammal and poultry IAV lineages indicating sustained respiratory transmission (***Figure 2C***). Conversely, the spectra of two minor NA lineages within domestic Anseriformes cluster with those from wild waterbird viruses (***Figure 2C***), implying that these lineages have retained a gastrointestinal transmission route, potentially explaining their low infection frequency in the population (***Figure S8***).

We thus conclude that gastrointestinal transmission results in exposure of viruses to (as yet undefined) shared mutagens that cause similar impacts on mutational spectra across wild waterbird and bat lineages, whereas sustained respiratory transmission exposes the virus to different airway-specific mutagens leading to distinct, convergent changes to mutational spectra across mammals, poultry, gulls, and domestic Anseriforme lineages.

### Detection of additional host-specific mutational signatures in IAV

We also identified several additional host-specific mutational signatures acting on IAV spectra that could potentially be used to infer infection source.

By comparing the spectrum changes between host jumps into mammals and those into gallinaceous poultry (***Figure S3***), we found that A>G is elevated in poultry while C>T is elevated in mammals (***Figure 3A***); the latter elevation only occurs in T[C>T] contexts (***Figure 3B***), a finding consistent with the action of APOBEC (apolipoprotein B mRNA editing enzyme, catalytic polypeptide) proteins which are known to favour this nucleotide context (*23*, *42*). Since T[C>T] mutations are reduced in poultry compared to both wild waterbirds and mammals (which exhibit similar levels, ***Figure 3B***), we hypothesised that this difference is driven by the loss of a mutagenic APOBEC protein in poultry. We examined the phylogenetic distribution of APOBEC genes across IAV hosts and found that APOBEC1 has been lost early within the Galliformes lineage, after divergence of Megapodiidae but prior to the common ancestor of known Galliformes hosts of IAV (***Figure 3C***). This suggests that avian APOBEC1 mutagenises IAV and that the observed reduction of T[C>T] mutations in poultry lineages is due to ancestral loss of this protein.

**Figure 3.**
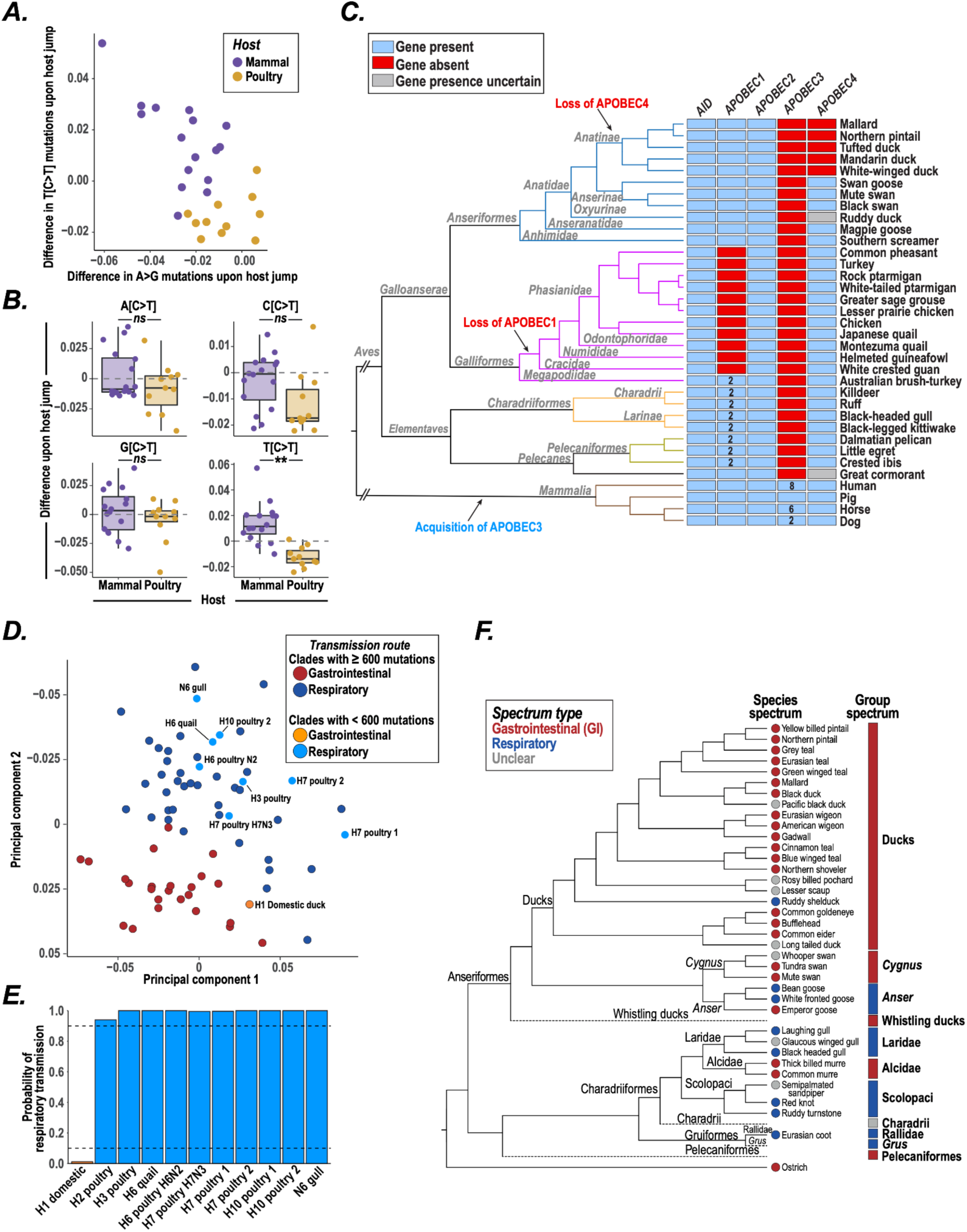
Calculation and application of IAV mutational signatures. (**A**) We compared changes in mutation type proportions upon host jumps into mammals and into poultry. Each point represents an individual mammal or poultry IAV lineage. (**B**) Comparison of changes in C>T mutations preceded by each possible nucleotide upon host jumps into mammals and into poultry. ** shows Benjamini Hochberg-corrected ANOVA *p* < 0.05; comparisons were carried out combining multiple lineages from the same host jump. (**C**) Presence and absence of individual APOBEC family genes across avian and mammalian species known to host IAV. Numbers within boxes show the number of gene copies for species with more than one copy. The cladogram shows evolutionary relationships between IAV hosts with available annotated genomes. Gene acquisition and loss events are inferred based on the distribution of presence and absence across the species cladogram. (**D-E**) We employed principal component analysis (**D**) and a likelihood-based classifier (**E**) to infer transmission route within ten IAV lineages with <600 mutations that have a clear predominant host but did not contain sufficient mutations to include in the main dataset. (**F**) We inferred predominant IAV infection site for individual avian species and higher taxonomic groupings by applying clustering (for spectra with >200 mutations) and likelihood-based classification to mutational spectra calculated on tip phylogenetic branches sampled from the corresponding species or taxonomic group. The cladogram represents phylogenetic relationships between species and groups.

By testing for differential signatures across the multiple independent origins of human and swine IAV lineages, we also found that C>A mutations were significantly elevated in human over swine lineages (ANOVA *p* < 0.05) and correspondingly detected a change in C>A mutations on each of three independent zoonotic and anthroponotic transmissions between these species (***Figure S9***).

Finally, by comparing mutational spectra of IAV across duck species calculated using mutations acquired on tip phylogenetic branches (see ***Methods***), we identified several instances of differential mutational patterns within isolates from individual species (such as black duck and American wigeon, ***Figure S10***).

### Inference of IAV transmission routes and infection sites from mutational spectrum

Due to the observed separation of IAV spectra by mode of transmission (***Figure 2C***), we next aimed to infer transmission routes from mutational spectra. To do this, we developed a likelihood-based classifier (see ***Methods***) that was able to correctly infer transmission route from mutational spectrum in 96% of cases (***Figure S11***), and applied this classifier together with mutational spectrum clustering to investigate transmission routes for small IAV lineages.

We first examined 11 minor IAV lineages, each with under 200 mutations, that are associated with specific host species (***Table S2***) and found strong support for sustained respiratory transmission of the gull lineage and nine poultry lineages (isolated from chickens and a variety of non-chicken Galliformes; ***Figure S12***) and gastrointestinal transmission of the domestic Anseriformes lineage (***Figures 3D-E***); findings consistent with those observed for larger IAV lineages from the same hosts (***Figure 2C***) and suggesting that reliable inferences can be achieved with low numbers of mutations.

While the spectra we describe above, calculated across IAV lineages, are dominated by mutations acquired on internal phylogenetic branches during transmission chains and can therefore be used to infer transmission route, the spectra calculated from tip phylogenetic branches include a higher proportion of mutations acquired during the infection period of the sampled individual, and can therefore reveal the major sites of the sampled infection. We therefore calculated the tip spectra of individual species and higher taxonomic groupings of wild waterbirds and found that IAV causes gastrointestinal infection in ducks, swans and alcids, but causes respiratory infections in gulls, shorebirds, *Gruiformes* and wild geese (after the divergence of the Emperor goose) (***Figures 3F***, ***S13***).

Sequences from respiratory wild waterbird groups are spread across subtypes (***Figure S14***), demonstrating that the respiratory association is not due to an individual viral lineage, but rather indicating that any IAV causes respiratory infection in these species. We identified eight HA sites that are enriched for mutations in tip branches from respiratory wild waterbird groups, including site 225 in the receptor binding domain (RBD) and other sites that may impact receptor binding and/or virus stability (***Figure S15***, ***Table S3***). However, tip branch spectra of the respiratory wild bird groups remain respiratory when excluding branches that acquire mutations at these sites (***Figure S15***). This strongly suggests that, while virus mutations may assist with respiratory tropism, IAV does not require any new mutations to cause respiratory infections in wild waterbirds. This supports host genetic differences (for example the distribution of sialic acid receptors (*43–45*)) being the major determinants of site of infection.

Since wild waterbird lineage spectra are gastrointestinal (***Figure 2C***) and alcids and swans are unlikely to drive IAV ecology (since the former is absent (*46*) and the latter present only at low densities (*47*) in freshwater environments ), our data supports ducks as being the major IAV reservoir, consistent with previous inferences based on detection rates (*48–51*). Conversely, respiratory infections are very unlikely to transmit onwards efficiently in wild birds, suggesting that gulls, geese, shorebirds and *Gruiformes* are unlikely to greatly contribute to IAV reservoirs or transmission dynamics (*52–54*), with the exception of HPAI GsGd and gull-adapted lineages.

We finally wondered whether we could apply our methodology to identify transmission routes driving IAV outbreaks within new species or populations. To test this, we calculated the mutational spectrum of a recent harbor seal H10N7 outbreak within the Weddell Sea (*55–58*). Despite this spectrum containing only 53 mutations, we found strong support that the outbreak was driven by sustained respiratory transmission (probability 0.996, ***Figure S16***, see ***Methods***). This suggests that mutational spectrum analysis can be applied in real time to resolve transmission routes (and, in some cases, species) driving outbreaks.

### Mutational spectra reveal the determinants of sustained respiratory transmission

We next sought to identify HA mutations associated with virus adaptation to sustained respiratory transmission by analysing mutations acquired around the transition from gastrointestinal to respiratory spectra in major lineages, identified through ancestral state reconstruction (***Figure 4A***, see ***Methods***). Due to the large number of mutations on these branches (199 sites mutate around at least one spectrum change; ***Table S4***, ***Figure S17***), we focussed on sites that mutate convergently across multiple spectrum changes and ascribed likely functional consequences of these changes from prior experimental analyses, including deep mutational scanning (DMS, see ***Methods***) (*59*).

**Figure 4.**
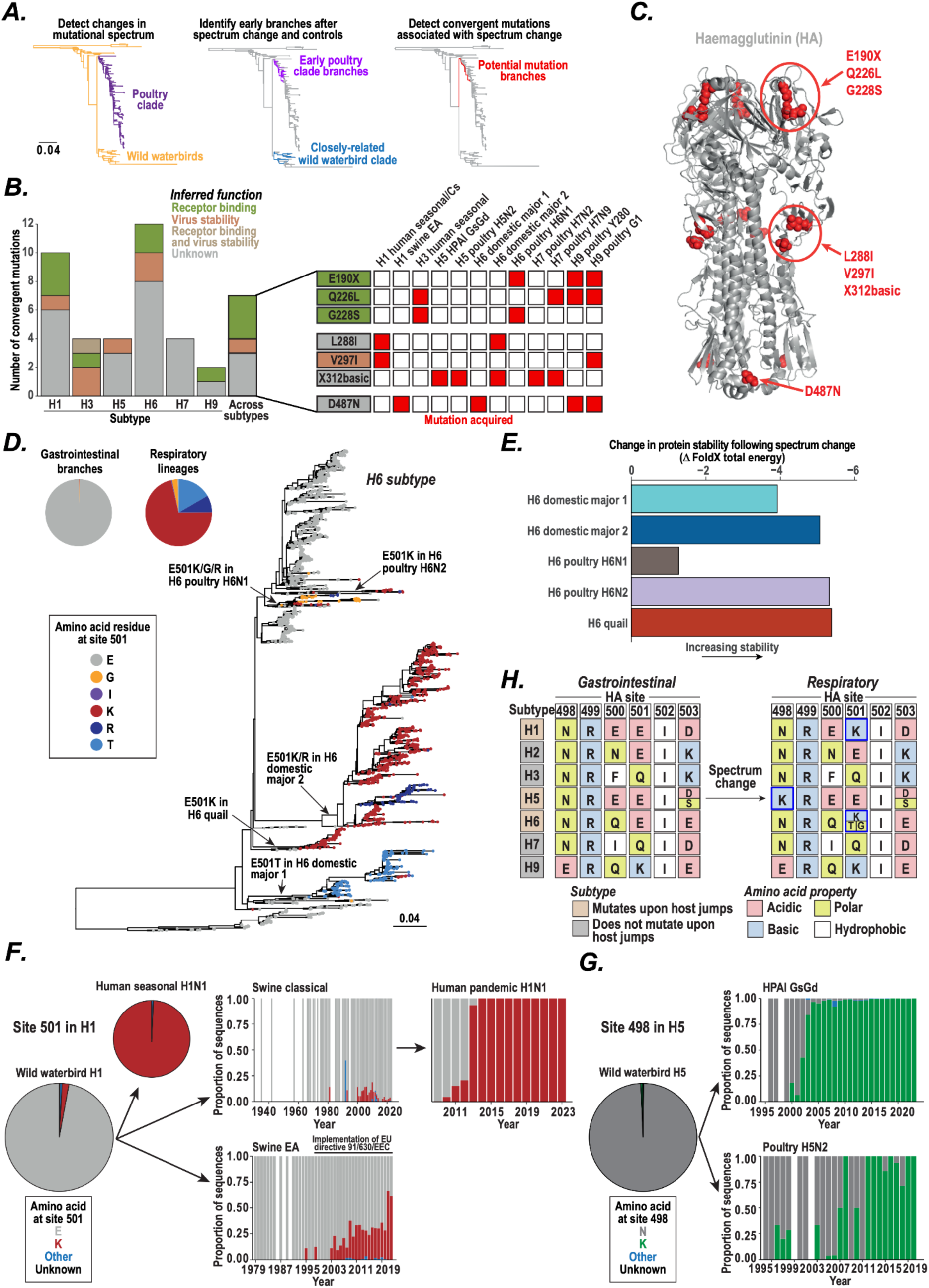
Identification of mutations contributing to sustained respiratory transmission. (**A**) To identify mutations that may contribute to tropism change, we calculated mutational spectra for 1) small groups of branches close to the root of individual sustained respiratory lineages and 2) for small groups of closely related branches outside of sustained respiratory lineages. This showed a gastrointestinal-like spectrum for the upstream branches and a respiratory spectrum for the early respiratory lineage branches. We can therefore identify mutations that may assist with the change in tropism as those acquired on branches between the upstream gastrointestinal branches and the early respiratory lineage. (**B**) We identified HA mutations acquired convergently leading to transitions to sustained respiratory transmission within each subtype and across subtypes, and inferred potential functions of these using DMS data (*59*). The inset shows whether each mutation that is convergent across subtypes is acquired in each sustained respiratory lineage. (**C**) The structural locations of the seven HA mutations acquired convergently by sustained respiratory lineages across subtypes are shown on PDB accession 4FNK. (**D**) The H6 phylogenetic tree is coloured by amino acid residue at HA site 501; mutations at site 501 leading to sustained respiratory lineages are labelled. The pie chart inset shows the distribution of amino acid residues at site 501 in the wild waterbird gastrointestinal sequences and in lineages with spectrum evidence of sustained respiratory transmission. (**E**) We employed FoldX to calculate the change in HA trimer structure when incorporating mutations at sites 63, 75 and 501 observed leading to lineages with spectrum evidence of respiratory transmission. Mutations at these sites are inferred to increase stability in each case. (**F**-**G**) Emergence of the (**F**) E501K and (**G**) N498K mutations in sustained respiratory lineages of H1 and H5, respectively. Pie charts show the distribution of amino acid residues at the respective site within all sequences from the respective lineage while bar charts show the distribution of amino acid residues at the respective site in sequences collected within each year. The years of implementation of EU directive 91/639/EEC introducing minimum space requirements for pigs is indicated above the plot for the swine EA lineage. (**H**) We identified the distribution of amino acid residues at each site surrounding site 501 in wild waterbird sequences from each subtype and in lineages with spectrum evidence of respiratory transmission from the subtype. We observe mutations that reduce acidity within the region of site 501 in subtypes that have an excess of acidic residues in this region.

We found seven sites that convergently mutate at the time of spectrum change across multiple IAV subtypes (***Figures 4B-C***, ***Table S5***). While three of these sites (190, 226 and 228, mature H3 site numbering used throughout (*59*, *60*)) cluster within the RBD (***Figures 4B-C***) and are known to impact receptor binding (*12*), the remaining four sites have not previously been associated with phenotypic impacts. Three of these four sites (288, 297 and 312) cluster together close to the HA1-HA2 boundary (***Figure 4C***) and will (based on DMS data) likely increase virus stability and alter receptor binding (***Figures 4B****, **S18***; ***Table S5***). The remaining substitution D487N, acquired in four spectrum changes, is located within the HA stalk domain and does not have a clear predicted phenotype.

We also identified 33 sites that mutate convergently leading to multiple spectrum changes within individual HA subtypes (***Figure 4B***, ***Table S5***). Some of these sites have well characterised phenotypes, for example G225D/E within subtype H1 impacts receptor binding (*12*). Between existing literature and DMS data, we could infer likely functions for 42% of these sites (***Table S5***), but phenotypic impacts for the remaining 58% should be the focus of future studies. Interestingly, the G146S substitution, important for airborne transmission of canine H3N2 by altering receptor binding (*7*), was acquired convergently by human seasonal H3N2 and equine H3N8 (***Table S5***), highlighting this as a key mutation for sustained respiratory transmission of H3.

We observed particularly striking convergent evolution in the branches leading to the five lineages within the H6 subtype with sustained respiratory transmission, identifying ten HA sites significantly associated with spectrum change (***Table S6***, see ***Methods***). Of these, sites 63, 75 and 501 appear to be particularly important as they mutate on branches leading to all, or almost all, spectrum shifts (***Figures 4D***, ***S19***; ***Table S6***). As these sites are unlikely to influence receptor binding (due to their structural location and lack of impact in DMS studies (*59*)), we hypothesised that they assist sustained respiratory transmission by increasing virus stability. To test this, we inserted the mutations at sites 63, 75 and 501 into the respective ancestral wild waterbird IAV genetic backgrounds and found, by structural modelling with *FoldX* (*61*), a large increase in stability in each case (***Figure 4E***), supporting a key role for these sites in enabling efficient respiratory transmission through increased virus stability.

We found that site 501, or the nearby site 498, was also mutated in branches leading to sustained respiratory lineages in H1 (where E501K was acquired leading to both human seasonal H1N1 and early within H1 2009 pandemic isolates, ***Figure 4F***) and H5 (where N498K expanded within HPAI GsGd and H5N2 poultry lineages, ***Figure 4G***). We observed that subtypes H1 (with the exception of swine lineages), H5, and H6 consistently undergo mutations that reduce negative charge around site 501 upon transitions to sustained respiratory transmission, while subtypes H2, H3, H7 and H9 do not, and wondered whether this difference may be driven by standing diversity in stability amongst wild waterbird viruses due to variation in amino acid residues close to site 501. We found that wild waterbird viruses within subtypes H1, H5 and H6 have an excess of acidic amino acids around site 501, while the remaining subtypes do not (***Figure 4H***). Together, these results suggest that: amino acid residues close to site 501 are key regulators of virus stability; excess acidic residues in this region reduces virus stability; and subtypes with excess acidic residues acquire mutations to reduce acidity to undergo sustained airborne transmission.

This raises the question of why E501K was not acquired early within the two swine H1 lineages. We hypothesise that E501K is not required in these lineages due to a predominance of direct contact transmission amongst swine (*6*) which removes the need for elevated stability to undergo efficient respiratory transmission. In support of this idea, we find that E501K has gradually increased in prevalence amongst Eurasian swine in the swine EA lineage since 2003, shortly after the implementation of European Union directive 91/630/EEC (***Figure 4F***); a regulation which introduced minimum space requirements for swine husbandry, thereby reducing rates of direct contact and therefore favouring airborne transmission and consequently the emergence of E501K to increase virus stability. In contrast, E501K has not expanded within the classical swine lineage within the USA, where no such regulations on swine farming have been introduced.

## Discussion

Our findings demonstrate the ability of mutational spectra to reveal the transmission dynamics and mechanisms of adaptation of IAV. We show that IAV mutational spectra are predominantly driven by transmission route, not by host phylogeny, causing IAV lineages that transmit through the same route to have similar mutational spectra, and implying that the (currently uncharacterised) niche-specific gastrointestinal and respiratory mutagens driving these mutational signatures are shared across species. Separately, we also identified additional host-associated mutational signatures that, for example, allow us to distinguish between human, poultry, and swine IAV lineages (despite them all undergoing sustained respiratory transmission), and could therefore be used to identify transmission routes and host species driving IAV outbreaks within new populations. Through mutational spectrum analysis of entire lineages and phylogenetic tip branches, we were able to characterise both the transmission route and the site of infection across IAV lineages infecting wild waterbirds, identifying ducks as the major reservoir for IAV. Our results therefore support the adoption of mutational spectra as an important surveillance tool to enable early detection of shifts in IAV transmission routes, thereby highlighting lineages more likely to undergo sustained transmission in humans.

By using shifts in mutational spectra, rather than host jumps, to identify critical evolutionary events, we were able to identify specific HA mutations across IAV subtypes that drive early adaptations enabling sustained respiratory transmission. Through this approach, we identified recurrent changes in multiple HA regions across IAV subtypes, revealed a convergent mutational process increasing virus stability, and highlighted the HA region neighbouring site 501 as a key determinant of sustained respiratory transmission for multiple IAV subtypes.

It is very likely that similar spectrum shifts exist in other microbes that infect multiple species and transmit through different routes. Our results therefore have immediate implications for surveillance and control of other human and zoonotic pathogens, and provide the first evidence that, in addition to revealing transmission routes and host species, mutational spectra can be applied to identify adaptive mutations and mechanisms.

## Acknowledgements

**Funding:** This work was supported by The Wellcome Trust (grant 107032AIA to R.A.F. and C.R.; grant 226602/Z/22/Z to R.A.F and C.R.; and grant 306920/Z/23/Z to T.S.); the Botnar Foundation (grant 6063 to R.A.F and C.R.); the UK Cystic Fibrosis Trust (Innovation Hub Grant 001 to R.A.F., J.P., and C.R.); the NIHR Cambridge Biomedical Research Centre (R.A.F. and C.R.); Coordenação de Aperfeiçoamento de Pessoal de Nível Superior (CAPES, to A.D.R.); and Conselho Nacional de Desenvolvimento Científico e Tecnológico (CNPq, to A.D.R.). **Authors contributions:** C.R. and R.A.F. conceived the project. C.R., T.P.P. and R.A.F. designed the experiments. C.R. and R.A.F. wrote the manuscript. C.R. and M.M. performed the phylogenetic analysis. C.R. performed the mutational spectrum analysis. C.R., T.S. and M.N. performed the likelihood analysis. A.D.R., V.M. and P.H.M.T. performed the computational structural modelling. R.A.F. and J.P. provided supervisory support. **Competing interests:** The authors declare no competing interests. **Data and materials availability:** all sequence accession numbers are listed in ***Table S7*** and ***Table S8***. All data are available in the manuscript, supplementary materials or at https://github.com/chrisruis/influenza_mutational_spectra.

## Supplementary Materials

Materials and Methods

Supplementary Text

Figs. S1 to S19

Tables S1 to S9

References (76–106)

