## Supplementary text and figures for "Mutational spectra reveal influenza virus transmission routes and adaptation"

### Materials and Methods

Calculation of mutational spectra

To calculate IAV single base substitution (SBS) mutational spectra, we first assembled sequencing datasets for each subtype and lineage using GISAID, NCBI GenBank and/or datasets in previous publications (see ***Supplementary text*** for details for each lineage). GISAID sequence accessions can be found in EPI_SET_251120bq (DOI: https://doi.org/10.55876/gis8.251120bq; ***Table S7***). NCBI GenBank accession numbers are included within ***Table S8***. We filtered sequences to remove those containing <90% of the complete HA or NA coding sequence for the respective subtype and those containing internal stop codons. Sequences were aligned at the amino acid level using MUSCLE (*62*) and alignments checked and adjusted manually in SeaView v5.0.4 (*63*). A maximum likelihood phylogenetic tree was reconstructed for each dataset using IQ-TREE v2.1.3 (*64*) employing the HKY model of nucleotide substitution with gamma rate heterogeneity and four gamma classes. Where necessary to identify initial lineages (see ***Supplementary text***), we reconstructed phylogenetic trees for large sequence datasets with FastTree v2.2 (*65*) employing the GTR model of nucleotide substitution; lineages were identified in these trees as outlined in the ***Supplementary text*** before reconstruction of a maximum likelihood phylogenetic tree for the lineage using IQ-TREE as above.

Phylogenetic trees were rooted either by maximising the heuristic residual mean squared value between sample isolation date and root-to-tip divergence using TempEst v1.5.3 (*66*) or by employing root locations defined in previous publications. In all cases, the root-to-tip correlation of the resulting trees was examined using TempEst and outlier sequences that have accumulated more or less diversity than expected given their collection date removed (*66*); where outliers were removed a new maximum likelihood phylogenetic tree was reconstructed using IQ-TREE as above and checked for the absence of outliers.

We examined the distribution of host species in detail across each phylogenetic tree, applying sequence names, associated metadata and original publications. Small phylogenetic clades predominantly containing sequences from poultry and/or domestic Anseriformes were separated from wild waterbird lineages; we applied MutTui v2.0.2 (*26*) to label the root ancestor and descendent branches of these clades, thereby splitting them into a separate spectrum.

We calculated mutational spectra using MutTui v2.0.2 (*26*), employing the sequence alignments and maximum likelihood phylogenetic trees outlined above. Where multiple lineages were analysed within a single phylogenetic tree, we applied MutTui to assign separate labels to phylogenetic branches within each lineage; MutTui thereby calculated an independent mutational spectrum for each lineage. Mutational spectra are presented in the genomic strand.

We included the 61 mutational spectra with at least 600 mutations within the main dataset. We chose this cutoff as we have previously shown that 600 mutations is sufficient to accurately represent a mutational spectrum (*26*). Mutational spectra containing fewer than 600 mutations but more than 200 mutations were used to test the likelihood-based classifier and spectrum clustering methods on spectra with fewer mutations.

Rescaling of mutational spectra

We rescaled mutational spectra to account for underlying sequence composition, which impacts the opportunity for different mutations to occur (*26*). As IAV undergoes frequent mutation, it is possible that the sequence composition can vary across a subtype or lineage such that the composition of a single reference sequence does not accurately represent the composition across the lineage. We therefore developed a new approach that we term "spectrum tree rescaling". This approach rescales a mutational spectrum taking into account the relative proportion of branch length that has been spent with a specific sequence composition across the phylogenetic tree. We calculate the composition (the frequency of each nucleotide triplet) at each node in the phylogenetic tree and multiply this composition by the sum of the lengths of the two immediately descending branches. We then sum the resulting scaled compositions across all nodes and rescale this to sum to one. This results in an overall reference composition for the phylogenetic tree. The lineage mutational spectrum is then rescaled by dividing each SBS mutation count by the frequency of its starting triplet within this lineage composition.

We compared symmetric mutation types using rescaled spectra. To identify symmetric mutation type pairs with a significant elevation of one mutation type over the other, we compared the median mutation ratio for each symmetric mutation type pair in the real data with the distribution of median mutation ratios in 1000 randomisations of mutation type proportions across the mutation types in the symmetric pair. The p-value was calculated as the proportion of randomisations with a ratio at least as large as with the real data (accounting for the direction of the ratio in the real data) and Benjamini-Hochberg correction applied. The mutation ratios were significant for C>A vs G>T, T>C vs G>A and T>G vs A>C. We report the median ratio across spectra as the fold enrichment for the symmetric pair.

<u>Calculation of synonymous mutational spectra</u>
To calculate synonymous mutational spectra, we ran MutTui as above but specifying the synonymous option. This infers the coding impact of all mutations on each phylogenetic branch and filters to only retain synonymous mutations. Note that where multiple mutations within the same codon occur on the same branch, their combined impact is applied.

<u>Analysis of the impact of surrounding nucleotide context</u>
We examined the impact of nucleotide context using rescaled SBS mutational spectra. We filtered each mutation type to retain lineage mutational spectra with at least 160 mutations of that mutation type. To determine whether there is significant variation in mutation frequency between contexts, we compared the variance across contexts within a mutation type in a given spectrum to the distribution of variances from 1000 random samplings of the observed number of mutations from a uniform distribution of contexts; the p-value was calculated as the number of random samplings with variance at least as large as in the real data and p-values were rescaled using the Benjamini-Hochberg method. Out of the 372 mutation types across spectra that were tested, 371 exhibited Benjamini-Hochberg-corrected p-values $< 0.05$, supporting strong context dependence across all mutation types. We identified outlier contexts mutated more or less often than expected as those whose median proportion across spectra (again filtering to retain only those with at least 160 mutations in the mutation type) is more than 2.5 times the median absolute deviation away from the median across all contexts. Potential ZAP contexts were identified as those that either introduce or remove a CG dinucleotide (*31*).

Comparison of mutational spectra

To identify changes in mutational spectrum upon host jumps, we subtracted mutation type proportions in the respective ancestral wild waterbird spectrum from each mammal or poultry spectrum (***Table S1***). We identified mutation types that change significantly upon host jumps by comparing the median difference for each mutation type upon host jumps in real data with the distribution of differences across 1000 randomisations of mutation type proportions within the mutation type across spectra. We included a single median value for all spectra that originate from the same host jump from wild waterbirds (***Table S1***). The p-value was calculated as the proportion of randomisations with a difference at least as large as in the real data and Benjamini-Hochberg correction was applied.

To examine differential mutational signatures between mammal and poultry spectra, we carried out an ANOVA comparing the changes in each mutation type across IAV jumps into mammals and into poultry from wild waterbirds. We compared the level of C>A mutations between human IAV lineages and swine IAV lineages through an ANOVA, run on median values from the same lineage. We identified mutation types elevated or reduced in wild waterbird species inferred to cause gastrointestinal infections as those more than 2.5 times the median absolute deviation away from the median proportion of the respective mutation type across species.

Temporal reconstruction of the H6 subtype

To identify introduction dates of H6 into domestic Anseriformes, we reconstructed a temporal phylogenetic tree. We assembled a dataset containing 400 H6 sequences, including 393 randomly sampled sequences and seven selected to represent the diversity close to the root of the two major domestic Anseriformes lineages. We reconstructed the temporal history of the lineage using BEAST v2.6.6 (*67*, *68*), employing the HKY model of nucleotide substitution. We used a relaxed log-normal clock model with a log-normal prior on the substitution rate with the mean set to 4.51E-3 (chosen as the slope of the RTT regression line of the H6 maximum likelihood tree analysed with TempEst) and standard deviation 0.5. We modelled the population history using a coalescent Bayesian skyline population prior. We carried out five independent runs and assessed convergence with Tracer v1.7 (*69*). The maximum clade credibility tree was identified using TreeAnnotator v2.6.4 (*70*).

Identification of APOBEC genes in IAV host species

To identify *APOBEC* genes, we searched annotations within available genomes for each taxonomic group that is known to host IAV using the NCBI Genome resource (genome accessions are listed in ***Table S9***). We further confirmed these annotations by aligning the annotated gene sequences. Where an *APOBEC* gene was not identified, we ran blastx of the corresponding *APOBEC* gene from the most closely related species with the gene to confirm that the gene was not present. If no BLAST hits were identified, we considered the gene to be absent. *APOBEC4* is not annotated in the genomes of ruddy duck or great cormorant; blastx resulted in a high identity match but only over a short region of the gene. We therefore treated the presence of *APOBEC4* in ruddy duck and great cormorant as uncertain.

Inference of IAV transmission route from mutational spectrum

We developed a likelihood-based classifier building on previous work to infer the likelihood that a set of SARS-CoV-2 mutations has been generated by molnupiravir exposure compared to typical

SARS-CoV-2 evolution (*21*). We calculate the likelihood of generating an observed set of mutations under two different mutational spectra; either the median wild waterbird spectrum vs the median mammal spectrum, or the median wild waterbird spectrum vs the median poultry spectrum (comparing across distinct dominant transmission routes in each case). We then calculate the likelihood ratio of these likelihoods and convert this into the probability that the observed mutations were generated by one spectrum over the other.

To test this classifier, we employed the dataset of known wild waterbird, mammal and poultry spectra. We removed each spectrum one at a time from the training dataset, calculated the median spectra excluding that spectrum and then identified the probability of that spectrum being generated by the known host spectrum. We considered a probability of at least 0.9 as being strong support that the mutation set was generated by the given spectrum.

Inference of infection sites for wild waterbird species and groups

We calculated mutational spectra for individual wild waterbird species and higher taxonomic grouping by combining mutations on tip phylogenetic branches sampled from the respective species or group. To identify infection sites, we examined clustering of each spectrum with more than 200 mutations within a principal component analysis including the individual spectrum and the 61 lineage dataset as in ***Figure 2C***. We additionally analysed all mutational spectra with at least 100 mutations through the likelihood-based classifier; we ran both wild waterbird vs poultry and wild waterbird vs mammal comparisons and assigned support to a transmission route if at least one of these comparisons resulted in a probability of at least 0.9. Where both comparisons resulted in intermediate probabilities, we assigned the infection site as unclear.

To identify HA sites where mutations are enriched on tip branches isolated from wild bird species and groups that exhibit a respiratory spectrum, we ran a Poisson test comparing the number of observed mutations on such tip branches with the number expected given the total tip branch length for respiratory species, the number of mutations at the site in duck tip branches and the total tip branch length leading to sequences isolated from ducks. We applied the Benjamini-Hochberg correction to p-values.

To show that enriched mutations are not required for respiratory infection, we calculated mutational spectra for wild bird groups with respiratory spectra excluding tip branches that acquire one or more of these mutations. We then applied the likelihood-based classifier as above and identified strong support for respiratory infection in each case (probability > 0.975).

Analysis of seal H10N7 outbreak lineage

We assembled a dataset containing all H10N7 HA sequences isolated from seals available on GISAID and a closely related H10N7 sequence isolated from poultry (*55–58*). We aligned the sequences and reconstructed a phylogenetic tree as above and rooted the tree on the poultry isolate. We calculated the mutational spectrum of the outbreak lineage using MutTui. To examine the dominant transmission route during the outbreak, we took two approaches: in the first, we applied the likelihood-based classifier comparing the likelihood of generating the 53 mutations acquired in the outbreak lineage under the median wild waterbird and median mammal spectrum; in the second, we compared the likelihood of generating the 53 mutations acquired in the outbreak lineage under the wild waterbird H10 spectrum and this spectrum adjusted by the median change in mutation type proportions upon host jumps of IAV from wild waterbirds to mammals (***Figure 2A***). In both cases, we found strong support (probability > 0.99) of mammal-like mutational patterns and thereby respiratory transmission.

To confirm that the likelihood-based classifier is capable of distinguishing respiratory and gastrointestinal spectra with 53 mutations, we carried out 1000 random subsamplings of each wild bird and mammal spectrum within the 61 lineage dataset and calculated the probability of each 53 mutation subsample being generated by the median mammal spectrum over the median wild waterbird spectrum. This showed a clear separation between mammal and wild waterbird subsamples and only mammalian spectrum subsamples are inferred to have a probability of mammal over wild waterbird at least as great as that with the seal outbreak lineage (0.996, ***Figure S16***).

Identifying and investigating mutations associated with spectrum shifts

In the above analyses, we identified lineages associated with a change in mutational spectrum from gastrointestinal to respiratory. To identify mutations associated with this spectrum shift, we first inferred phylogenetic branches on which the spectrum shift may have occurred. To do this, we calculated the mutational spectrum of the early internal branches within sustained respiratory lineages and confirmed that this spectrum is respiratory through spectrum clustering and the likelihood-based classifier. This confirms that the shift to a respiratory spectrum occurred within these early branches or earlier. We additionally calculated the mutational spectrum of a clade of wild waterbird sequences that clusters close to the sustained respiratory lineage and confirmed that this spectrum is gastrointestinal through spectrum clustering and the likelihood-based classifier. This confirms that the shift to a respiratory spectrum occurred after the divergence between this wild waterbird clade and the early branches in the sustained respiratory lineage. This results in a small number of phylogenetic branches on which the spectrum shift may have occurred and therefore a small number of branches on which adaptive mutations important for this shift must have been acquired. Three lineages (human seasonal H1N1/H1 classical swine; H5 poultry H5N2; H9 G1) diverge into multiple large subclades at their root; we tested both subclades by spectrum clustering and found that they exhibited a respiratory-like spectrum in each case. This strongly supported the transition to respiratory spectrum being acquired by the root of these lineages, enabling us to further narrow down branches with potential mutations of interest to those preceding the lineage root. We identified amino acid substitutions on phylogenetic branches of interest by employing ancestral state reconstruction through treetime v0.8.1 (*71*). We were not able to examine H2 human H2N2 or canine H3N2 with this method due to insufficient mutations when splitting the lineage into early branches, or examine H3 equine H3N8 due to an intermediate probability for transmission route within the early branches.

We converted amino acid sites within each subtype to site numbering within the mature H3 protein using published site conversions (*59*, *60*), validated through comparison of our sequence alignments across subtypes. Convergent mutations were identified as the same H3 site mutating leading to multiple spectrum shifts.

We developed a Poisson test to identify mutations that are associated with spectrum shifts in the H6 subtype. We compared the observed number of mutations on branches leading to spectrum shifts with the expected number given the total branch length leading to spectrum shifts, the total branch length in the remainder of the tree and the number of mutations at the site across the remainder of the tree. We carried out Benjamini-Hochberg correction of the resulting p-values and identified ten sites associated with spectrum change (***Table S6***).

To ascribe likely functional consequences, we employed published literature describing functions of individual sites and mutations where possible, and applied DMS data including impacts on specific functions (*59*). We set cutoffs for functional impacts in DMS data using the minimum phenotypic impact of mutations known to influence phenotype. For receptor binding, we used sites 137, 190, 193, 224, 225, 226 and 228 (*59*). For virus stability, we used mutations Y17H, A19T, H24Q, E31K, H110Y and T318I (*59*, *72–74*). Sites exhibiting at least one mutation with an impact at least as great as the minimum of these sites were assigned the corresponding phenotype.

We mapped mutations onto existing protein structures to examine their structural location. To examine the impact of mutations at sites 63, 75 and 501 on virus stability within the H6 subtype, we applied structures generated through AlphaFold 3 (*75*). We initially validated the accuracy of AlphaFold 3 structural prediction for influenza HA by predicting the structure of a homotrimer of the amino acid sequence of Protein Data Bank (PDB) accession 6HJQ, which is a solved structure of the full length HA. The root mean square deviation (RMSD) between the solved and AlphaFold 3 predicted structures was 2.219 Angstroms, showing a highly accurate structural prediction. We therefore generated homotrimeric structures for the amino acid sequence at the phylogenetic node immediately preceding each spectrum change in H6 using AlphaFold 3. We incorporated into all three monomers of the respective ancestral virus the mutations at sites 63, 75 and 501 that each H6 lineage exhibiting a respiratory spectrum acquires (***Figures 4D***, ***S19***), and inferred their impact on protein stability using FoldX 5.1.

**Data availability**

Accession numbers for GenBank sequences are provided in ***Table S8***. GISAID sequence accessions can be found in ***Table S7*** and in EPI_SET_251120bq (DOI: https://doi.org/10.55876/gis8.251120bq). Data and code underlying the analyses can be found at https://github.com/chrisruis/influenza_mutational_spectra.

### Supplementary text

#### H1 subtype

Within the H1 subtype, we calculated mutational spectra for the full diversity of H1 in wild waterbirds (spectrum H1_wild_waterbird), human seasonal H1N1 pre-2009 (spectrum H1_human_seasonal_H1N1), human pandemic H1N1 (H1pdm09, spectrum H1_human_pandemic_H1N1), H1N1 classical swine (spectrum H1_swine_Cs) and H1N1 swine EA (Eurasian avian, spectrum H1_swine_EA) (*76*). Human seasonal H1N1 and H1 classical swine form a cluster that diverges from the avian sequences at the root of the tree (*77*). It is unclear whether one of the human seasonal H1N1 and H1 swine classical lineages descends from the other, or whether they are the result of independent spillovers from wild waterbirds (*77*). We therefore treated these lineages as a single spillover for statistical tests. H1 human pandemic H1N1 (H1pdm09) descends from H1 classical swine (*78*) so is also included within this spillover for statistical tests. H1 swine EA descends from an independent spillover event and clusters within the diversity amongst wild waterbird H1 sequences (*76*, *79–81*); swine EA was therefore treated as a separate spillover in statistical tests.

To calculate the H1 wild waterbird spectrum, we downloaded all available H1 avian sequences from GISAID and reconstructed a phylogenetic tree containing these sequences and representatives from close to the root of each of the mammalian H1 clades, removing several wild waterbird sequences that clustered within human or swine lineages. The remaining sequences form a single monophyletic clade that clusters as a sister lineage to human seasonal H1N1, classical swine and human pandemic H1N1, as expected from previous studies (*77*, *78*). We separated a small domestic Anseriformes lineage sampled in ducks in China from 2011-2017 (spectrum H1_domestic). We calculated the H1 wild waterbird mutational spectrum using all mutations across the remaining avian clade.

To calculate the human seasonal H1N1 mutational spectrum, we downloaded all available sequences within the "Seasonal H1N(1) type" from GISAID. We removed several outliers based on root-to-tip (RTT) distance and lab-passaged isolates. The tree was rooted based on previous studies (*78*) which provided a strong RTT correlation with no remaining outliers.

To calculate the human pandemic H1N1 (H1pdm09) mutational spectrum, we used the dataset of sequences applied in the Nextstrain (*82*) H1pdm09 "pandemic" and "12y" datasets, which covers the period of circulation of this lineage. We downloaded the corresponding sequences from GISAID and NCBI GenBank. There were no outliers in this dataset based on RTT distance.

To examine the swine lineages, we initially reconstructed a phylogenetic tree of all swine H1 sequences available on GISAID (n = 15,108) using FastTree (*65*) (see ***Methods***). From this, we extracted H1 classical swine sequences and swine EA sequences as those clustering within the previously designated clade 1A and clade 1C (*83*), respectively, applying sequences close to the root of each clade based on previous publications (*78*, *81*, *83*).

To calculate the H1 classical swine mutational spectrum, we analysed four designated clades (*83*) independently (clade 1A.1.1, clade 1A.2, clade 1A.3.3.2 and clade 1A.3.3.3) and combined the resulting mutational spectra. Each clade was extracted from the above FastTree based on clade designations in Anderson et al 2016 (*83*). A maximum likelihood tree was reconstructed for each clade using IQ-TREE (*64*) (see ***Methods***) and outliers removed based on RTT.

To calculate the swine EA mutational spectrum, we extracted sequences in clade 1C in the above FastTree containing all swine H1 sequences. We reconstructed a maximum likelihood tree of these sequences using IQ-TREE (*64*) and removed several outliers based on RTT distance.

**H2 subtype**

Within the H2 subtype, we calculated mutational spectra for the full diversity of H2 isolated from wild waterbirds (spectrum H2_wild_waterbird) and for the human H2N2 pandemic lineage (spectrum H2_human_pandemic_H2N2) which circulated amongst humans between 1957 and 1968 (*84*).

To calculate the H2 human pandemic H2N2 mutational spectrum, we assembled a dataset containing all H2 sequences on GISAID and extracted the human H2H2 lineage from a phylogenetic tree reconstructed on this dataset and validated that there were no outliers based on RTT distance.

To calculate the H2 wild waterbird mutational spectrum, we removed the human sequences from the complete H2 dataset and reconstructed a phylogenetic tree on the remaining avian sequences with IQ-TREE. There were no outliers based on RTT distance. We separated a small poultry-associated lineage sampled in the USA from 1990-1997 (spectrum H2_poultry).

**H3 subtype**

Within the H3 subtype, we calculated mutational spectra for the complete diversity of H3 isolated from wild waterbirds (spectrum H3_wild_waterbird) and for three mammalian lineages: human seasonal H3N2 (which has circulated continuously amongst humans since 1968 (*84*), spectrum H3_human_seasonal_H3N2), equine H3N8 (which has circulated continuously in horses since at least 1963 (*85–87*), spectrum H3_equine_H3N8) and canine H3 (which has circulated continuously within dogs since around 2006 (*7*, *88*), spectrum H3_canine).

To calculate the H3 wild waterbird mutational spectrum, we downloaded all avian H3 sequences from GISAID and reconstructed a phylogenetic tree with FastTree containing these sequences and a subset of early seasonal human H3N2 sequences, a subset of equine H3N8 sequences and all available swine H3 sequences. We excluded avian sequences that clustered within the human seasonal H3N2 lineage. We reconstructed a phylogenetic tree of the remaining avian H3 isolates with IQ-TREE and removed several outlier sequences based on RTT distance. We separated a poultry-associated clade isolated in China from 2014-2022 (spectrum H3_poultry).

To calculate the H3 human seasonal H3N2 mutational spectrum, we utilised the mutational spectrum we calculated previously (*20*) based on a published H3N2 dataset (*89*). As the common ancestor of sequences in this dataset occurred in roughly 1998 (*89*), 30 years after the initial emergence of H3N2 into humans (*84*), we identified mutations close to the root of the human lineage using all available human H3N2 sequences available on GISAID sampled before 2000 (n = 1097); this includes 21 sequences collected before 1970 that cluster around the root of the clade.

To calculate the H3 equine H3N8 mutational spectrum, we downloaded all equine H3 sequences on GISAID and removed two outlier sequences based on phylogenetic clustering. A maximum likelihood phylogenetic tree of the remaining sequences was rooted on the earliest sequences based on RTT correlation with no outliers.

To calculate the H3 canine mutational spectrum, we downloaded all canine H3 sequences available on GISAID and reconstructed a phylogenetic tree with FastTree containing these sequences and all H3 sequences from wild waterbirds and a subset of human H3N2 sequences. This demonstrated that all of the canine sequences formed a single cluster with no spillover from dogs back into wild waterbirds, thereby confirming a single introduction into dogs. We therefore reconstructed a phylogenetic tree of the canine H3 sequences and removed several outlier sequences based on RTT distance.

**H4 subtype**

Within the H4 subtype, we calculated the mutational spectrum of the complete diversity of H4 isolated from wild waterbirds (spectrum H4_wild_waterbird). We downloaded all H4 sequences from GISAID and removed several outliers based on RTT distance. We did not identify any subclades within the H4 diversity associated with poultry or domestic Anseriformes; the wild waterbird mutational spectrum was therefore calculated on all branches of the phylogenetic tree.

**H5 subtype**

Within the H5 subtype, we calculated mutational spectrum for the diversity of H5 in wild waterbirds (spectrum H5_wild_waterbird), the H5 HPAI GsGd lineage (*9*) (spectrum H5_HPAI_GsGd) and a low pathogenic H5N2 lineage that has circulated within chickens since 1994 (*90–92*) (spectrum H5_poultry_H5N2).

We initially reconstructed a phylogenetic tree with FastTree containing all available H5 sequences on GISAID (n = 19047), within which we identified the HPAI GsGd and poultry H5N2 lineages. We identified the HPAI GsGd lineage as all sequences that clustered downstream of sequence A/goose/Guangdong/1/1996. There was a very strong RTT correlation within this lineage in the FastTree phylogenetic tree, with the exception of 7 outlier sequences that were removed. Due to the very large number of sequences within the HPAI GsGd lineage (n = 17351), we calculated the mutational spectrum of this lineage using a random sample of 2000 sequences sampled from wild waterbirds, on which a phylogenetic tree was reconstructed using IQ-TREE.

To examine the mutational spectrum of distinct host groups within HPAI GsGd (wild waterbirds, mammals and poultry; ***Figure S6***), we used the phylogenetic tree reconstructed with FastTree above. We labelled tip phylogenetic branches of the different host groups using MutTui (*26*) based on sequence names and metadata and calculated independent mutational spectra of these tip branch groups.

To calculate the mutational spectrum of H5 in wild waterbirds, we examined two large H5 wild waterbird clades visible within the above FastTree that were sampled over multiple decades. Phylogenetic trees for each clade were reconstructed with IQ-TREE and outliers based on RTT correlation removed. Mutational spectra were calculated for each clade independently and combined.

To calculate the H5 poultry H5N2 mutational spectrum, we extracted sequences within the corresponding clade of the phylogenetic tree reconstructed above with FastTree and reconstructed a phylogenetic tree using IQ-TREE. Several sequences were removed based on RTT distance.

**H6 subtype**

Within the H6 subtype, we reconstructed mutational spectra for the complete diversity of H6 in wild waterbirds (spectrum H6_wild_waterbird), two large domestic Anseriformes lineages (previously referred to as group I and group II (*93–96*), spectra H6_domestic_major_1 and H6_domestic_major_2) and a low pathogenic H6N1 lineage that has circulated in poultry for multiple decades (*97*, *98*) (spectrum H6_poultry_H6N1).

We calculated the mutational spectrum of each of these lineages from a single phylogenetic tree, reconstructed with IQ-TREE on all H6 sequences available on GISAID (n = 3017) following removal of outlier sequences based on RTT distance. We labelled the phylogenetic tree with MutTui to separate the two major domestic Anseriformes lineages, the poultry H6N1 lineage and two additional poultry lineages (not included in the 61 lineage dataset due to containing <600 sampled mutations; these lineages are referred to as H6_poultry_H6N2 and H6_quail); the remaining branches were assigned to H6 wild waterbirds.

**H7 subtype**

Within the H7 subtype, we calculated mutational spectra for H7 in wild waterbirds (spectrum H7_wild_waterbird), a low pathogenic H7N2 lineage that has circulated in poultry for multiple decades (*99*) (spectrum H7_poultry_H7N2) and a H7N9 lineage that has undergone sustained transmission within poultry (*100*, *101*) (spectrum H7_poultry_H7N9).

We initially reconstructed a phylogenetic tree with FastTree of all H7 sequences available on GISAID (n = 5976). Within this tree, there were two large clades that diverge from the root when the tree is midpoint rooted. We reconstructed a phylogenetic tree for each of these two large clades independently with IQ-TREE and removed outlier sequences based on RTT distance. We reconstructed mutational spectra on these two clades independently. The poultry H7N2 lineage and poultry H7N9 lineage were separated through labelling with MutTui and we also separated three additional small poultry lineages (spectra H7_poultry_H7N3, H7_poultry_1 and H7_poultry_2). The remaining branches were assigned to the wild waterbird mutational spectrum and the two clade wild waterbird spectra were combined to generate the final H7 wild waterbird spectrum. The poultry H7N9 lineage contains a large number of human samples; we excluded tip phylogenetic branches leading to human samples from the poultry H7N9 mutational spectrum.

**H8 subtype**

Within the H8 subtype, we calculated the mutational spectrum of the complete diversity of H8 isolated from wild waterbirds (spectrum H8_wild_waterbird). We downloaded all H8 sequences from GISAID and removed one outlier based on RTT distance. We did not identify any subclades within the H8 diversity associated with poultry or domestic Anseriformes; the wild waterbird mutational spectrum was therefore calculated on all branches of the phylogenetic tree.

**H9 subtype**

Within the H9 subtype, we calculated mutational spectra of the complete diversity of H9 isolated from wild waterbirds (spectrum H9_wild_waterbird), the low pathogenic poultry lineage Y280 (35) (spectrum H9_poultry_Y280) and the low pathogenic poultry lineage G1 (35) (spectrum H9_poultry_G1).

We initially reconstructed a phylogenetic tree with IQ-TREE of all H9 sequences available on GISAID (n = 13729). Within this tree, there were several large clades predominantly isolated from poultry that included the Y280 and G1 lineages (*35*). We reconstructed independent phylogenetic trees for each of these lineages and for the remaining wild waterbird sequences using IQ-TREE and removed outliers based on RTT distance.

Due to the large number of sequences within the Y280 lineage (n = 10953), we calculated the Y280 mutational spectrum employing a phylogenetic tree reconstructed with IQ-TREE and containing a random sample of 2000 Y280 sequences.

We did not identify any subclades within the remaining H9 wild waterbird diversity associated with poultry or domestic Anseriformes; the wild waterbird mutational spectrum was therefore calculated on all branches of the phylogenetic tree.

**H10 subtype**
Within the H10 subtype, we calculated the mutational spectrum of the complete diversity of H10 isolated from wild waterbirds (spectrum H10_wild_waterbird). We downloaded all H10 sequences from GISAID and removed several outliers based on RTT distance. We separated two poultry lineages (spectra H10_poultry_1 and H10_poultry_2) and calculated the wild waterbird spectrum across the remaining phylogenetic branches.

**H11 subtype**
Within the H11 subtype, we calculated the mutational spectrum of the complete diversity of H11 isolated from wild waterbirds (spectrum H11_wild_waterbird). We downloaded all H11 sequences from GISAID and removed several outliers based on RTT distance. We did not identify any subclades within the H11 diversity associated with poultry or domestic Anseriformes; the wild waterbird mutational spectrum was therefore calculated on all branches of the phylogenetic tree.

**H12 subtype**
Within the H12 subtype, we calculated the mutational spectrum of the complete diversity of H12 isolated from wild waterbirds (spectrum H12_wild_waterbird). We downloaded all H12 sequences from GISAID; no outliers were identified based on RTT distance. We did not identify any subclades within the H11 diversity associated with poultry or domestic Anseriformes; the wild waterbird mutational spectrum was therefore calculated on all branches of the phylogenetic tree.

**H13 subtype**
The H13 subtype is associated with infection of gulls (27). We downloaded all H13 sequences from GISAID; no outliers were identified based on RTT distance. Within an H13 phylogenetic tree reconstructed with IQ-TREE, almost all sequences were isolated from gulls with no evidence of sustained transmission outside of the Laridae family. We therefore calculated the mutational spectrum of the complete diversity of H13 (spectrum H13_gull).

**H16 subtype**

The H16 subtype is associated with infection of gulls (*27*). We downloaded all H16 sequences from GISAID; no outliers were identified based on RTT distance. Within an H16 phylogenetic tree reconstructed with IQ-TREE, almost all sequences were isolated from gulls with no evidence of sustained transmission outside of the Laridae family. We therefore calculated the mutational spectrum of the complete diversity of H16 (spectrum H16_gull).

**N1 subtype**

Within the N1 subtype, we calculated mutational spectra for the full diversity of N1 in wild waterbirds (spectrum N1_wild_waterbird), human seasonal H1N1 (spectrum N1_human_seasonal_H1N1), human pandemic H1N1 (H1pdm09, spectrum N1_human_pandemic_H1N1), H1N1 classical swine (spectrum N1_swine_Cs), H1N1 swine EA (Eurasian avian, spectrum N1_swine_EA) and an N1 lineage associated with the H5 HPAI GsGd lineage (*9*) (spectrum N1_HPAI_GsGd).

Human seasonal H1N1 and H1N1 classical swine form a cluster (77); it is unclear whether one of the human seasonal H1N1 and H1N1 swine classical lineages descends from the other, or whether they are the result of independent spillovers from wild waterbirds (*77*). We therefore treated these lineages as a single spillover for statistical tests. H1N1 swine EA descends from an independent spillover event and clusters within the diversity amongst wild waterbird N1 sequences (*76*, *79–81*). The N1 of human pandemic H1N1 (H1pdm09) descends from H1N1 classical swine (*78*). We therefore treated the N1 from human pandemic H1N1 and N1 from H1N1 swine EA as a single spillover in statistical tests.

To calculate the mutational spectra of the wild waterbird and HPAI GsGd lineages, we initially reconstructed a phylogenetic tree with FastTree containing all avian N1 sequences on GISAID with a sample of early diversity from human seasonal H1N1, human pandemic H1N1, H1N1 classical swine and H1N1 swine EA; we removed several avian sequences that clustered within the mammalian lineages. Furthermore, within this FastTree there is a clear lineage of NA sequences whose corresponding HA is within the H5 HPAI GsGd lineage (*9*). We separated sequences within the HPAI GsGd lineage from the remainder of the avian sequences and reconstructed independent phylogenetic trees with IQ-TREE to calculate the mutational spectra of the N1 HPAI GsGd lineage and N1 in wild waterbirds, respectively. We did not identify any subclades within the N1 wild waterbird diversity associated with poultry or domestic Anseriformes; the wild waterbird mutational spectrum was therefore calculated on all branches of the phylogenetic tree.

To calculate the mutational spectrum of the human seasonal H1N1 lineage, we downloaded NA sequences within the "Seasonal H1N(1) type" dataset on GISAID. We reconstructed a phylogenetic tree of these sequences using IQ-TREE and removed several outliers based on RTT distance.

To calculate the human pandemic H1N1 (H1pdm09) mutational spectrum, we used the dataset of sequences applied in the Nextstrain (*82*) H1pdm09 "pandemic" and "12y" datasets, which covers the period of circulation of this lineage. We downloaded the corresponding sequences from GISAID and NCBI GenBank. There were no outliers in this dataset based on RTT distance.

To examine the swine N1 lineages, we initially reconstructed a phylogenetic tree using FastTree containing all swine N1 sequences available on GISAID (n = 8538), a sample of early diversity from the human seasonal H1N1 lineage and a sample of early diversity from the human pandemic H1N1 lineage. We extracted sequences within the N1 swine classical lineage (which clustered close to the human seasonal H1N1 lineage as expected (*78*)) and N1 swine EA lineage from this phylogenetic tree. We excluded sequences that cluster downstream of the root of the H1N1 human pandemic lineage. We reconstructed phylogenetic trees for each swine lineage independently using IQ-TREE and removed several outlier sequences based on RTT distance.

**N2 subtype**

Within the N2 subtype, we calculated mutational spectra for the complete diversity of N2 in wild waterbirds (spectrum N2_wild_waterbird), the poultry lineage Y280 (spectrum N2_poultry_Y280), the poultry lineage G1 (spectrum N2_poultry_G1), a low pathogenic poultry H5N2 lineage that has circulated within chickens since 1994 (*90–92*) (spectrum N2_poultry_H5N2), a low pathogenic poultry H7N2 lineage that has circulated in poultry for multiple decades (*99*) (spectrum N2_poultry_H7N2), one major and one minor lineage associated with domestic Anseriformes (spectra N2_domestic_major and N2_domestic_minor, respectively), human seasonal H3N2 (spectrum N2_human_seasonal_H3N2) and two swine N2 lineages termed N2.1998 and N2.2002 that descend from human seasonal H3N2 and have circulated continuously in swine for multiple decades (*102–105*) (spectra N2_swine_1998 and N2_swine_2002, respectively). As the swine N2.1998 and swine N2.2002 lineages descend from N2 human seasonal H3N2, we combined these three lineages for statistical tests. The Y280, G1, poultry H5N2 and poultry H7N2 lineages were combined with their corresponding HA lineages for statistical tests.

To calculate the N2 human seasonal H3N2 mutational spectrum, we used all human N3N2 NA sequences available on GISAID collected before 2000. We reconstructed a phylogenetic tree of these sequences using IQ-TREE and removed several outliers based on RTT distance.

To calculate the swine N2 lineage spectra, we initially reconstructed a phylogenetic tree with FastTree containing all swine N2 sequences available on GISAID (n = 12016) with a sample of sequences from early in human seasonal H3N2. We identified the roots of the swine N2.1998 and swine N2.2002 lineages based on previous publications (*102*). Due to the large number of sequences within the N2.2002 lineage (n = 7151), we calculated the swine N2.2002 mutational spectrum using a phylogenetic tree reconstructed using IQ-TREE on a random subset of 2000 sequences. We reconstructed a phylogenetic tree for the N2.1998 dataset with IQ-TREE and removed several outliers based on RTT distance.

To examine the avian lineages, we initially reconstructed a phylogenetic tree with FastTree containing a random sample of 3000 avian N2 sequences on GISAID, with a sample of early diversity from the N2 human pandemic H2N2, N2 human seasonal H3N2, swine N2.1998 and swine N2.2002 lineages. We excluded several avian sequences that clustered downstream of the mammalian lineages. Amongst the remaining sequences, there were several large clades sampled from poultry that corresponded to Y280, G1 the poultry H5N2 and poultry H7N2 lineages. We extracted the sequences within these clades and calculated mutational spectra using independent phylogenetic trees reconstructed with IQ-TREE.

We extracted the remaining avian sequences, which form the wild waterbird lineage, and reconstructed a phylogenetic tree with IQ-TREE. We separated the two domestic Anseriformes lineages through labelling with MutTui and calculated the mutational spectrum of the wild waterbird lineage using the remaining branches.

**N3 subtype**

Within the N3 subtype, we calculated mutational spectra for the complete diversity of N3 in wild waterbirds (spectrum N3_wild_waterbird), a minor domestic Anseriformes lineage (spectrum N3_domestic_minor) and a gull lineage (spectrum N3_gull).

We reconstructed a phylogenetic tree with IQ-TREE containing all N3 sequences on GISAID and removed several outlier sequences based on RTT distance. Within this phylogenetic tree, there is a clear gull clade that diverges from the root of the tree (linked with the H16 gull lineage) and a clade isolated from domestic Anseriformes. We separated these lineages through labelling with MutTui and calculated the wild waterbird mutational spectrum across the remaining phylogenetic branches.

**N4 subtype**

Within the N4 subtype, we calculated the mutational spectrum of the complete diversity of N4 isolated from wild waterbirds (spectrum N4_wild_waterbird). We downloaded all N4 sequences from GISAID and removed several outlier sequences based on RTT distance. We excluded small clades associated with the H5 HPAI GsGd lineage and with poultry, each of which contained a very small number of mutations. We calculated the N4 wild waterbird mutational spectrum across the remaining phylogenetic branches.

**N5 subtype**

Within the N5 subtype, we calculated the mutational spectrum of the complete diversity of N5 isolated from wild waterbirds (spectrum N5_wild_waterbird). We downloaded all N5 sequences from GISAID and removed several outlier sequences based on RTT distance. We excluded several small clades associated with the H5 HPAI GsGd lineage, each of which contained a very small number of mutations. We calculated the N5 wild waterbird mutational spectrum across the remaining phylogenetic branches.

**N6 subtype**

Within the N6 subtype, we calculated the mutational spectrum of the complete diversity of N6 isolated from wild waterbirds (spectrum N6_wild_waterbird), a large lineage associated with the H5 HPAI GsGd lineage (spectrum N6_HPAI_GsGd) and a major lineage in domestic Anseriformes (spectrum N6_domestic_major).

We initially reconstructed a phylogenetic tree with FastTree containing all N6 sequences available on GISAID (n = 6382). This showed two large clades that diverged from the root of the tree when midpoint rooted. We extracted the sequences within each of these two subclades and reconstructed independent phylogenetic trees with IQ-TREE. We removed several outlier sequences based on RTT distance. We separated a lineage associated with the H5 HPAI GsGd lineage, a large domestic Anseriformes lineage and a small N6 gull lineage (spectrum N6_gull). The N6 wild waterbird spectrum was calculated across the remaining phylogenetic branches; we combined the mutational spectra from the two subclades.

**N7 subtype**

Within the N7 subtype, we calculated the mutational spectrum of the complete diversity of N7 isolated from wild waterbirds (spectrum N7_wild_waterbird). We downloaded all N7 sequences from GISAID reconstructed a phylogenetic tree with IQ-TREE. This tree diverged into three clades close to the root: the equine H7N7 lineage and two avian lineages. We excluded the equine H7N7 sequences and analysed the two avian lineages. We reconstructed an independent phylogenetic tree for each lineage with IQ-TREE and removed several outlier sequences based on RTT distance. We excluded a small clade associated with poultry which contained a very small number of mutations and calculated the wild waterbird mutational spectrum across the remaining phylogenetic branches, combining the spectra calculated on the two clades.

**N8 subtype**

Within the N8 subtype, we calculated mutational spectra for N8 isolated from wild waterbirds (spectrum N8_wild_waterbird), a lineage associated with the H5 HPAI GsGd lineage (spectrum N8_HPAI_GsGd) and the equine H3N8 lineage which has circulated continuously in horse since at least 1963 (*85–87*) (spectrum N8_equine_H3N8).

We initially reconstructed a phylogenetic tree with FastTree of all N8 sequences available on GISAID (n = 7256). We extracted from this tree sequences that clustered within the equine H3N8 lineage, sequences that clustered within a lineage associated with the H5 HPAI GsGd lineage and a large North American wild waterbird lineage with a strong RTT correlation. We calculated mutational spectra for these lineages using independent phylogenetic trees reconstructed with IQ-TREE. A poultry-associated clade with a very small number of mutations was excluded from the wild waterbird lineage; the N8 wild waterbird mutational spectrum was calculated on the remaining phylogenetic branches.

**N9 subtype**

Within the N9 subtype, we calculated mutational spectra for the complete diversity of N9 isolated from wild waterbirds (spectrum N9_wild_waterbird) and a H7N9 lineage that has undergone sustained transmission within poultry (*100*, *101*) (spectrum N9_poultry_H7N9). The poultry H7N9 lineage was combined with its corresponding HA lineage for statistical tests.

We reconstructed a phylogenetic tree of all available N9 sequences on GISAID using IQ-TREE and removed several outlier sequences based on RTT distance. We excluded a small poultry-associate clade that contains very few mutations. We labelled the N9 poultry H7N9 lineage using MutTui and calculated the N9 wild waterbird spectrum across the remaining phylogenetic branches. We excluded tip branches leading to sequences isolated from humans from the N9 poultry H7N9 mutational spectrum.

### Supplementary figures

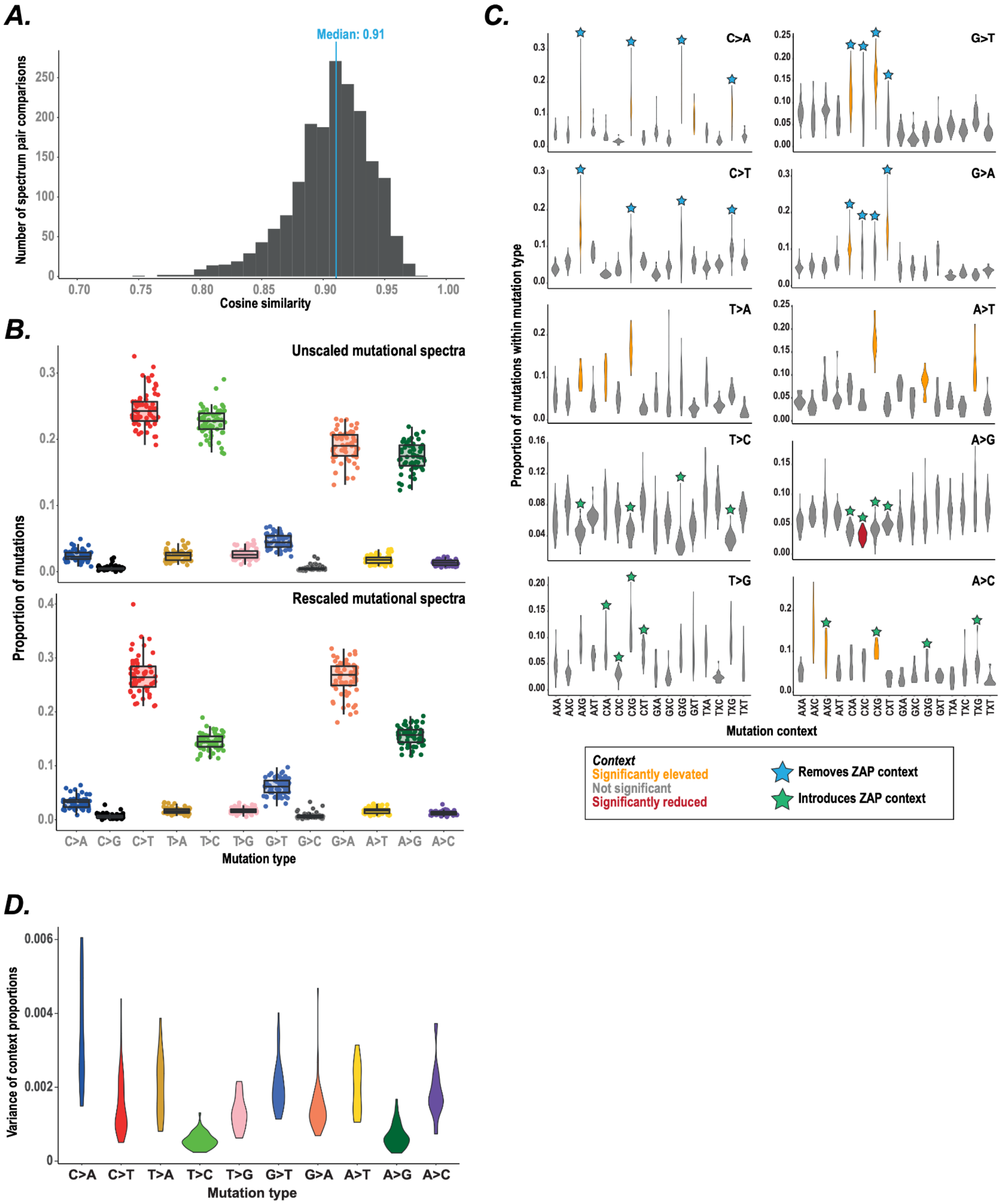


**Figure S1. Specific mutational patterns during IAV evolution.** (**A**) We calculated the cosine similarity between all pairs of SBS spectra across the 61 lineage dataset. (**B**) The proportion of each mutation type is shown for each lineage in unscaled mutational spectra (top panel) and in mutational spectra rescaled by sequence composition across the respective phylogenetic tree (bottom panel). (**C**) For each mutation type, we filtered to include lineages where the respective mutation type has at least 160 mutations. The proportion of mutations within the respective mutation type that occur within each context is shown. Mutations that introduce or remove CG dinucleotides and therefore ZAP contexts are indicated. Mutation contexts were identified as significantly elevated or reduced if their median proportion is more than 2.5 times the median absolute deviation away from the median context proportion for the mutation type. (**D**) We calculated the variance between SBS contexts for each mutation type within each lineage spectrum. The distribution of variances across lineage spectra is shown for each mutation type.

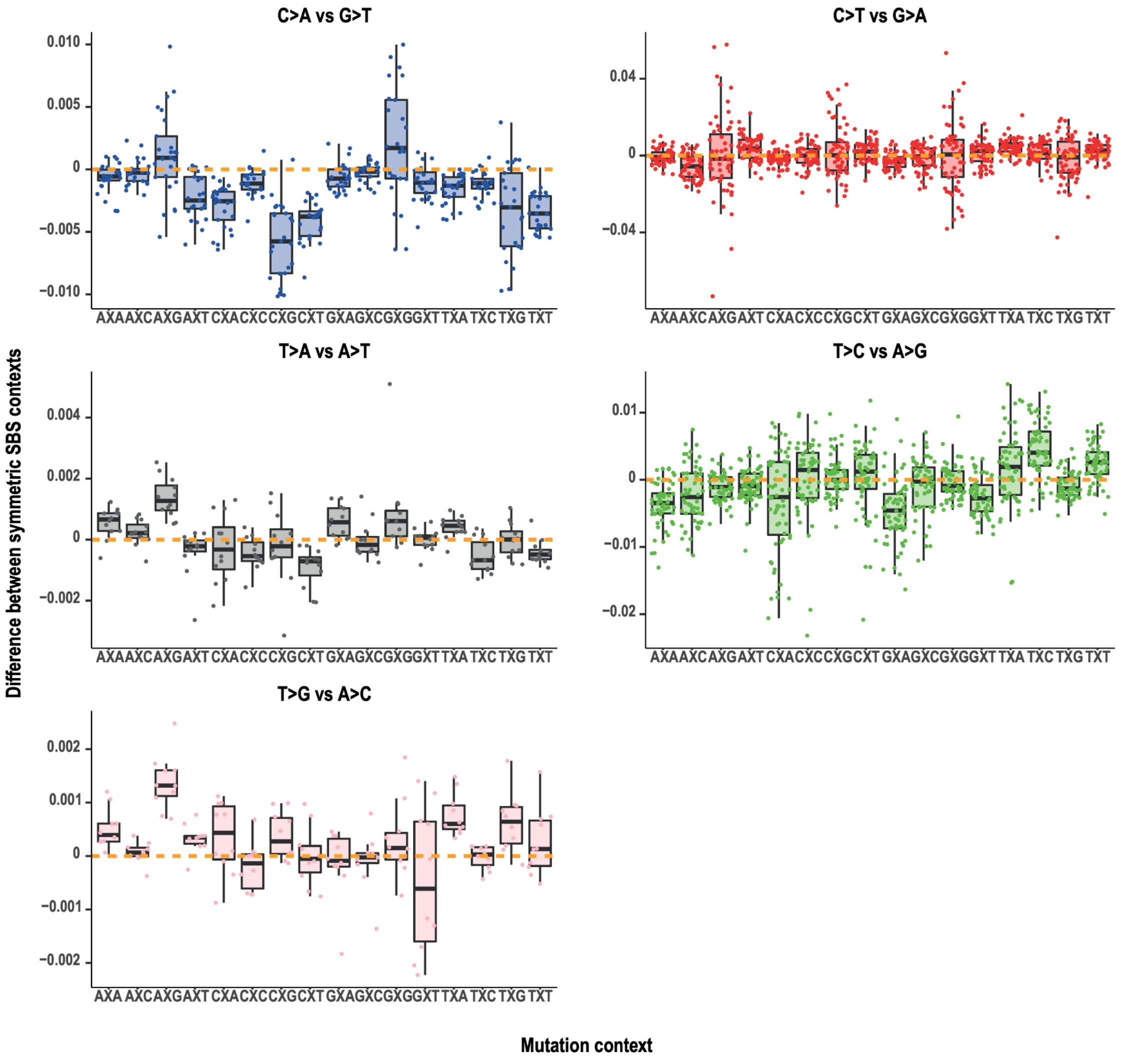


**Figure S2. Comparison of symmetric SBS contexts.** For each comparison, we filtered to keep lineage mutational spectra containing at least 160 mutations within each mutation type. We compared the frequency of symmetric SBS mutation type pairs across rescaled mutational spectra. The labelled context is that in the first named mutation type (i.e. C>A, C>T, T>A, T>C or T>G).

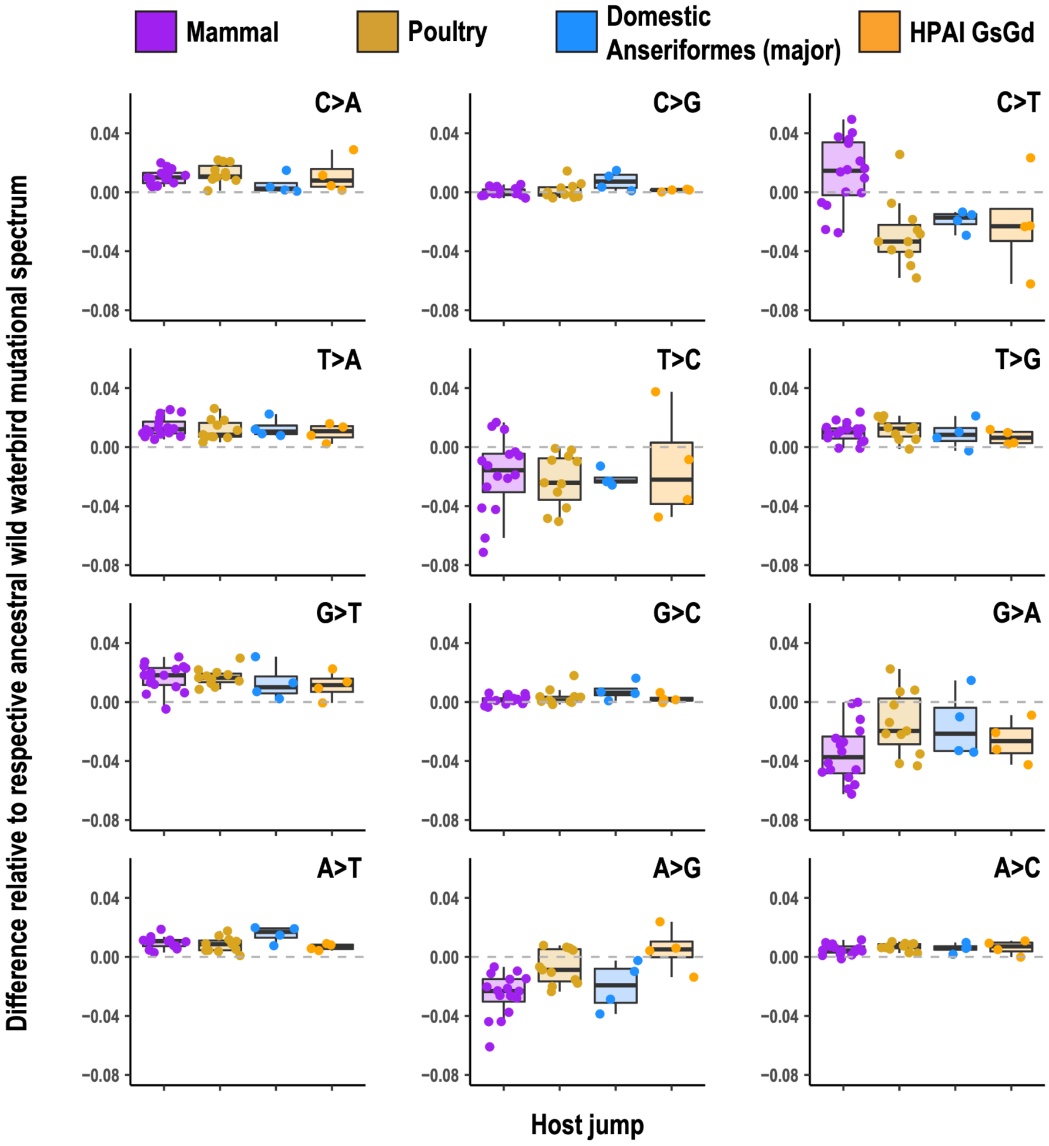


**Figure S3. Changes in mutational spectra upon IAV host jumps.** We identified mutational spectrum changes associated with host jumps by subtracting the respective ancestral wild waterbird mutational spectrum from each mammal, poultry, major domestic Anseriformes, and HPAI GsGd lineage spectrum.

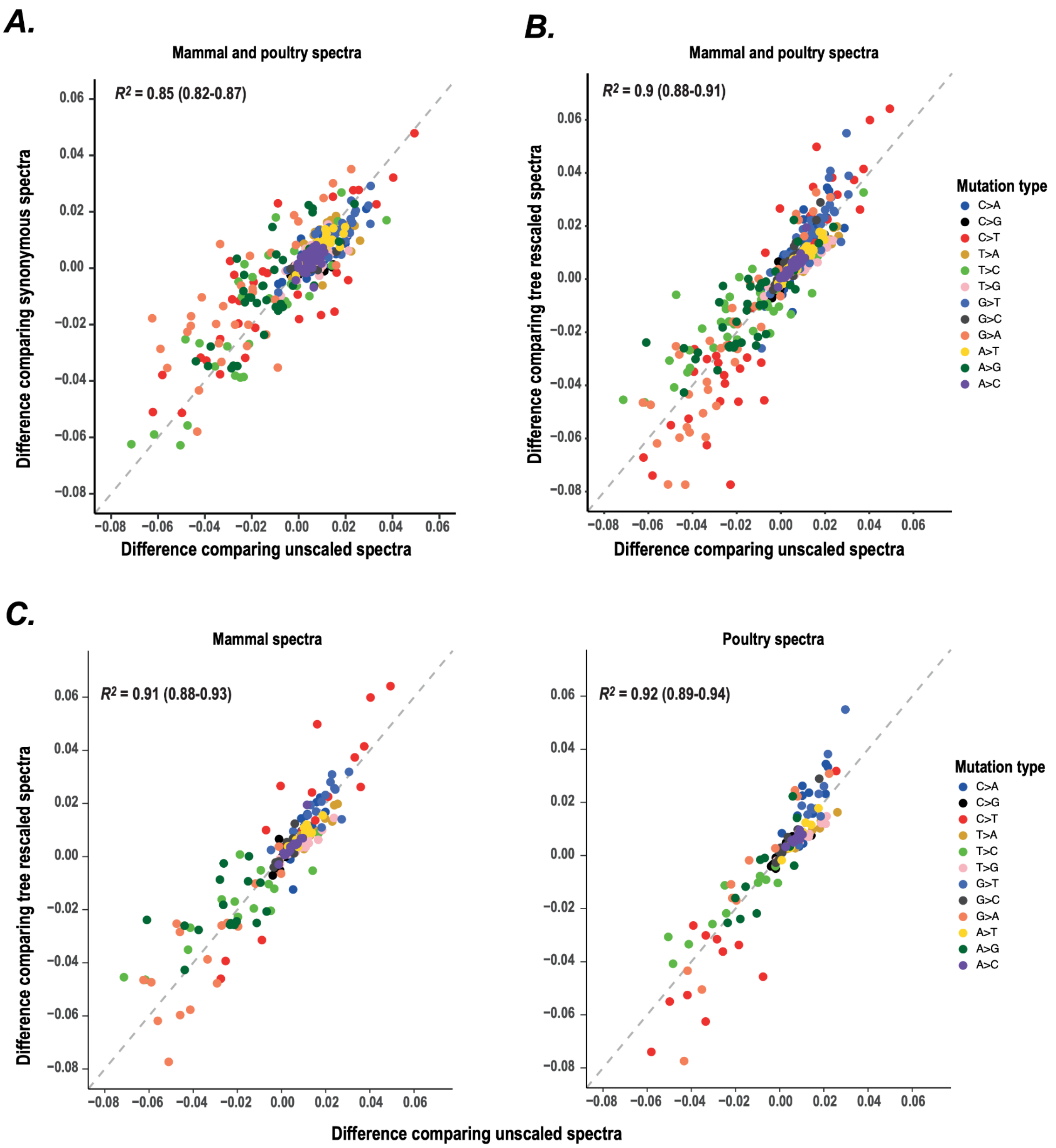


**Figure S4. Mutational spectrum changes upon host jumps are highly similar in unscaled, rescaled and synonymous mutation-only mutational spectra.** The proportion change for each mutation type upon host jumps is shown for (**A**) mammal and poultry lineages comparing unscaled spectra and synonymous mutation-only spectra, (**B**) mammal and poultry lineages comparing unscaled and rescaled spectra, (**C**) separated mammal and poultry lineages compared unscaled and rescaled spectra.

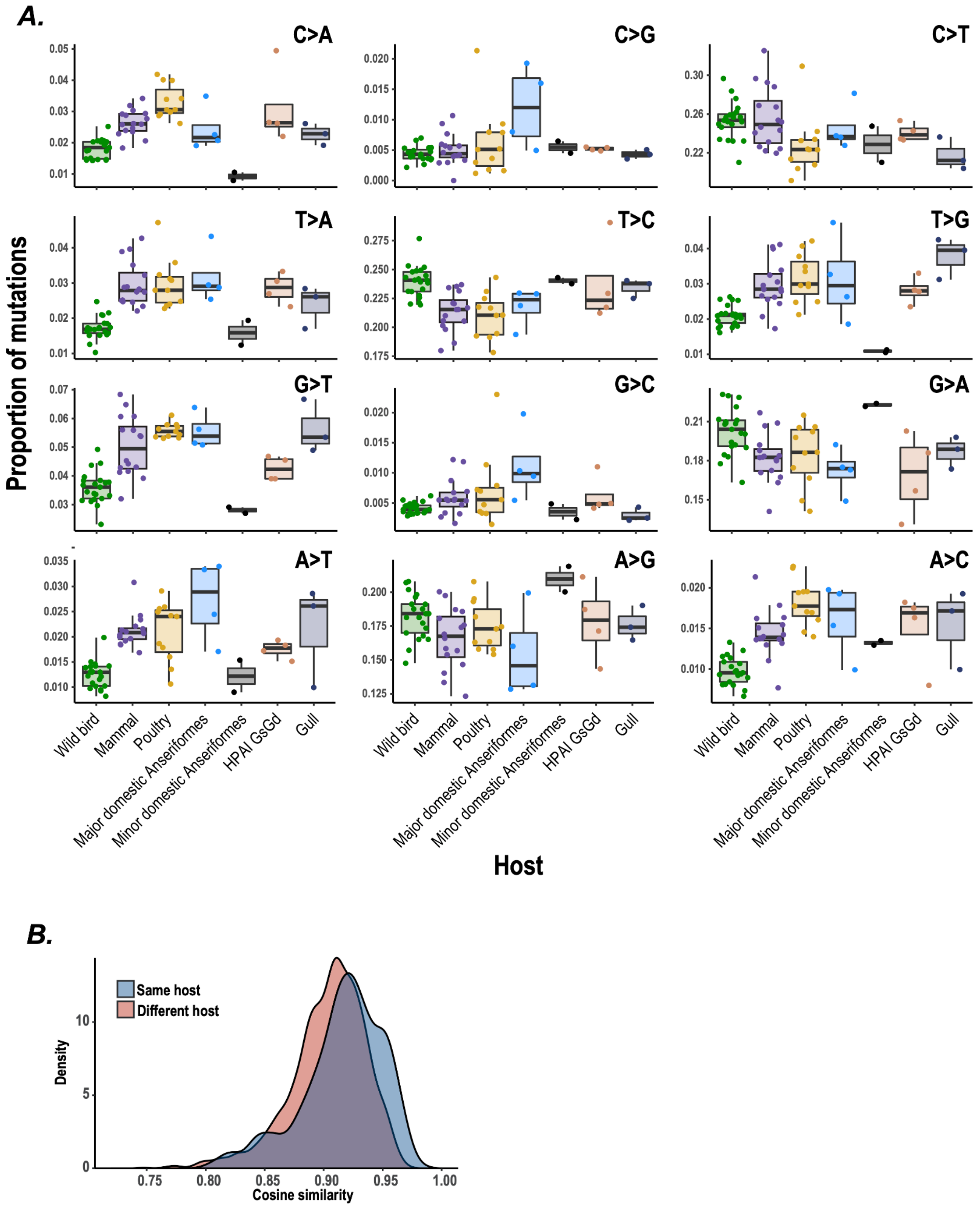


**Figure S5. IAV mutational patterns differ by host species.** (**A**) The proportion of each mutation type in unscaled mutational spectra is shown divided by host group. (**B**) The distribution of cosine similarity values between all pairs of SBS mutational spectra is shown divided by whether the compared lineage pair are from the same host group or a different host group.

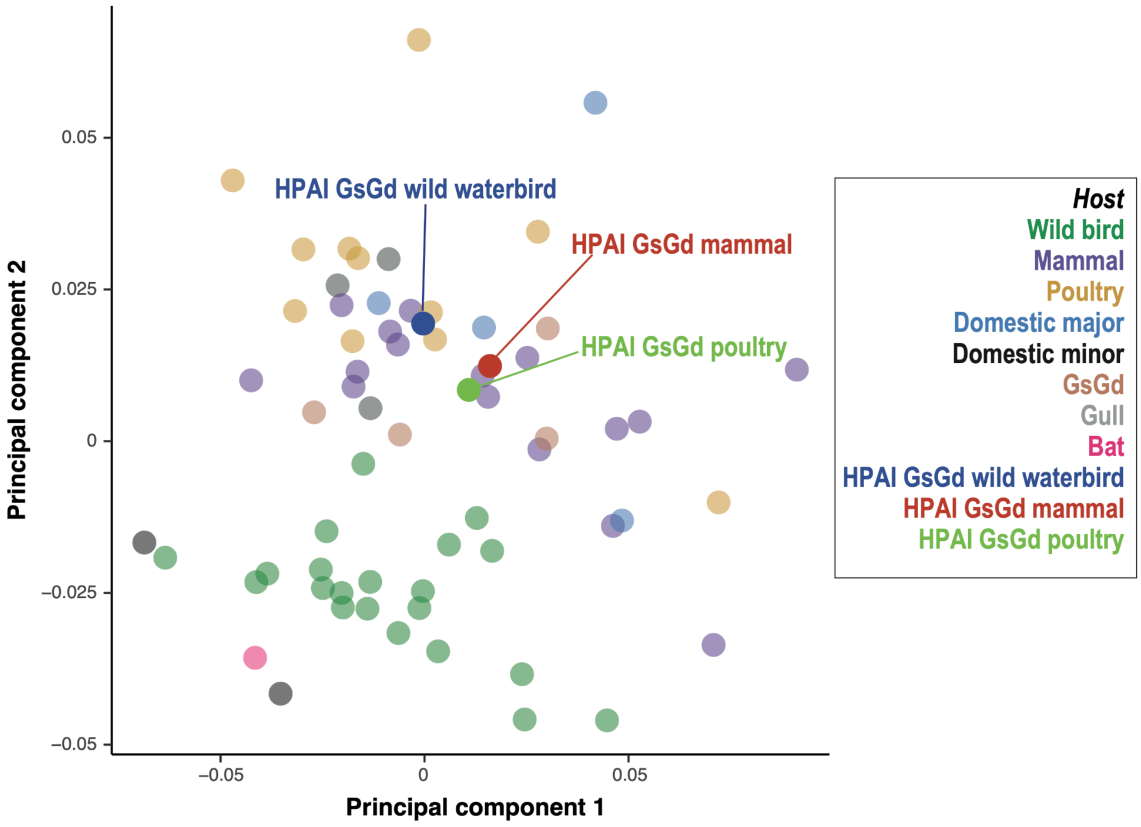


**Figure S6. The HPAI GsGd mutational spectrum is respiratory-like regardless of host.** We calculated mutational spectra for HPAI GsGd infecting different host groups by incorporating mutations acquired on tip phylogenetic branches isolated from the respective host group. Principal component analysis of the HPAI GsGd host-resolved mutational spectra with the 61 lineage dataset (as in ***Figure 2C***) shows that HPAI GsGd clusters with the mutational spectra of sustained respiratory lineages regardless of host.

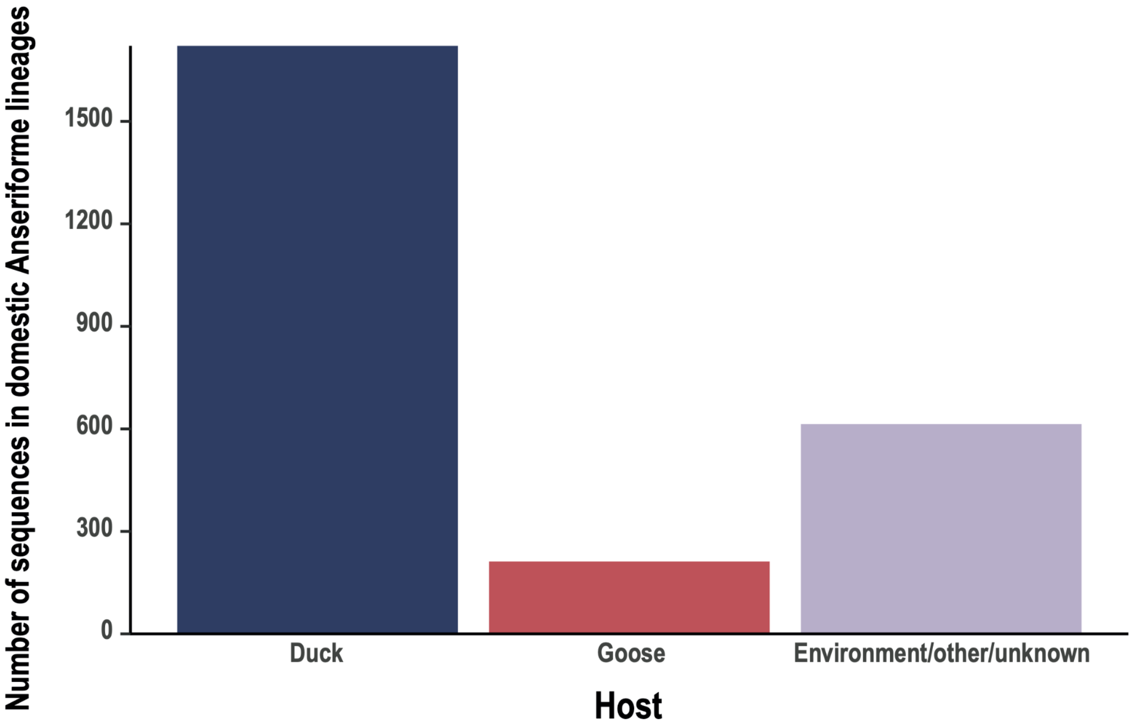


**Figure S7. Domestic Anserformes lineages are predominantly sampled from ducks.** We identified hosts for all sequences that cluster within a major or minor domestic Anseriformes lineage based on sequence names and associated metadata. The majority of the sequences are from domestic ducks with smaller numbers from domestic geese.

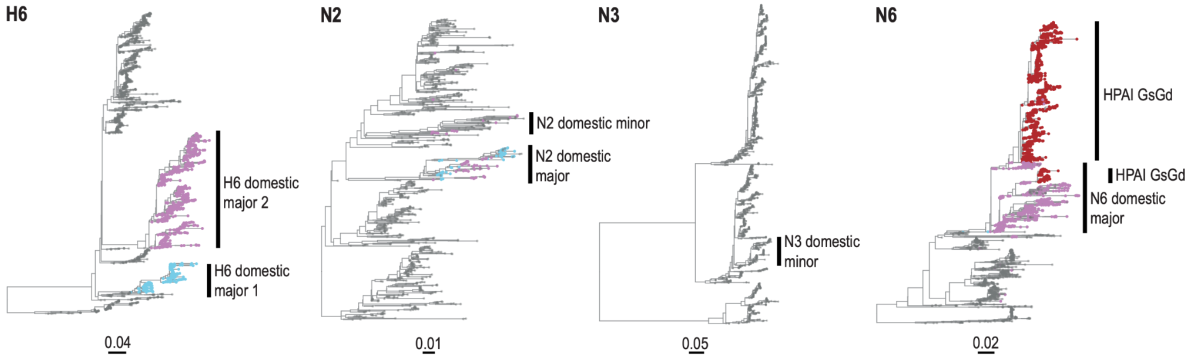


**Figure S8. Classification and linkage of domestic Anseriforme lineages.** Phylogenetic trees for IAV subtypes containing domestic Anseriforme lineages from the 61 lineage dataset are shown. The two major H6 domestic Anseriforme lineages and their corresponding NA sequences are labelled; these lineages were designated as major domestic Anseriforme lineages due to their high prevalence. The two NA lineages unlinked to the H6 lineages were designated as minor domestic Anseriforme lineages.

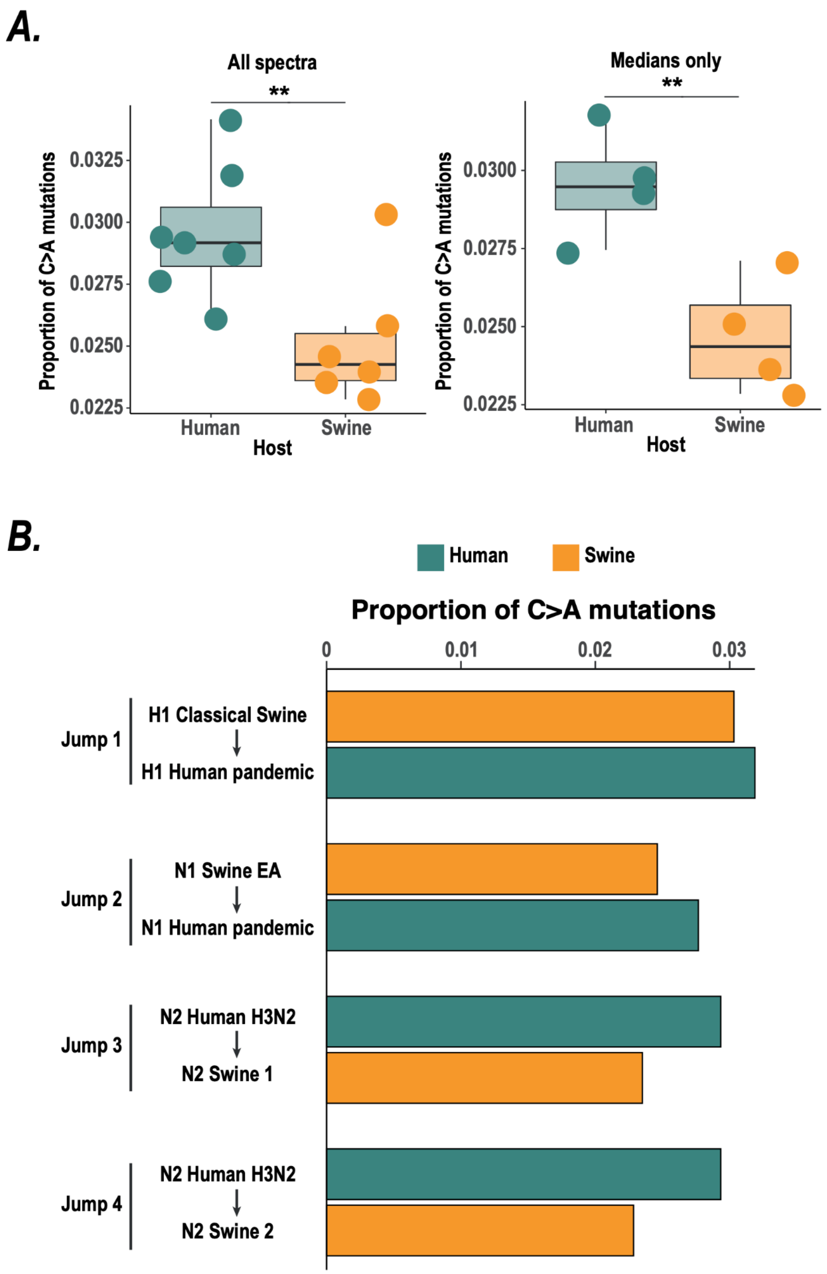


**Figure S9. IAV exhibits elevated C>A mutations in humans compared to swine.** (**A**) The left panel shows the proportion of C>A mutations for all human and swine IAV lineages. The right panel shows the median proportions of C>A mutations for each host jump into humans or swine, combining lineages that descend from the same host jump. ** shows ANOVA p-value < 0.05. C>A is significantly elevated in human IAV lineages compared to swine lineages in both comparisons. (**B**) The proportion of C>A mutations is shown across each jump of IAV between human and swine. C>A proportion increases upon jumps into humans and reduces upon jumps into swine.

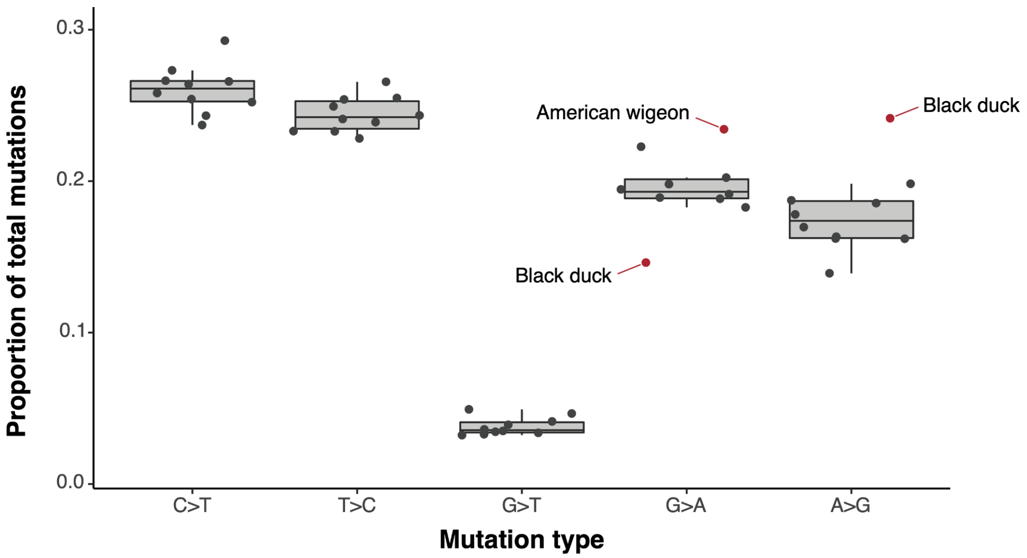


**Figure S10. Mutational patterns differ between IAV infections of wild waterbird species.** We calculated mutational spectra for individual species of wild waterbirds by combining mutations acquired on tip phylogenetic branches leading to the corresponding species (see ***Methods***). We compared the proportion of mutations belonging to each mutation type across wild waterbird species that exhibit gastrointestinal infections (***Figure 3F***). Mutation types that are elevated or reduced within a specific species (defined as the mutation type proportion being more than 2.5 median absolute deviations away from the median of the mutation type) are labelled in red.

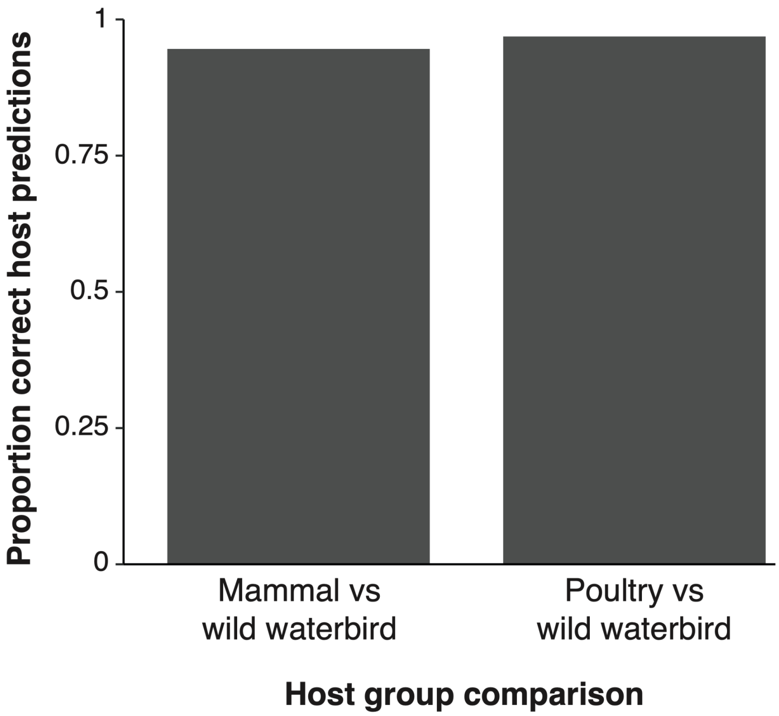


**Figure S11. Transmission route can be inferred from mutational spectrum through a likelihood-based classifier.** We developed a likelihood-based classifier to infer whether a set of mutations is best explained by either of two underlying mutational spectra (see ***Methods***). We applied this classifier to two comparisons, each across species groups with different predominant transmission routes. To test the ability of this classifier to correctly infer transmission route, we left each individual lineage spectrum out in turn, trained the classifier on the medians of the remaining lineage spectra and calculated the probability of the correct transmission route. The proportion of tests that resulted in a probability of the correct transmission route > 0.9 is shown.

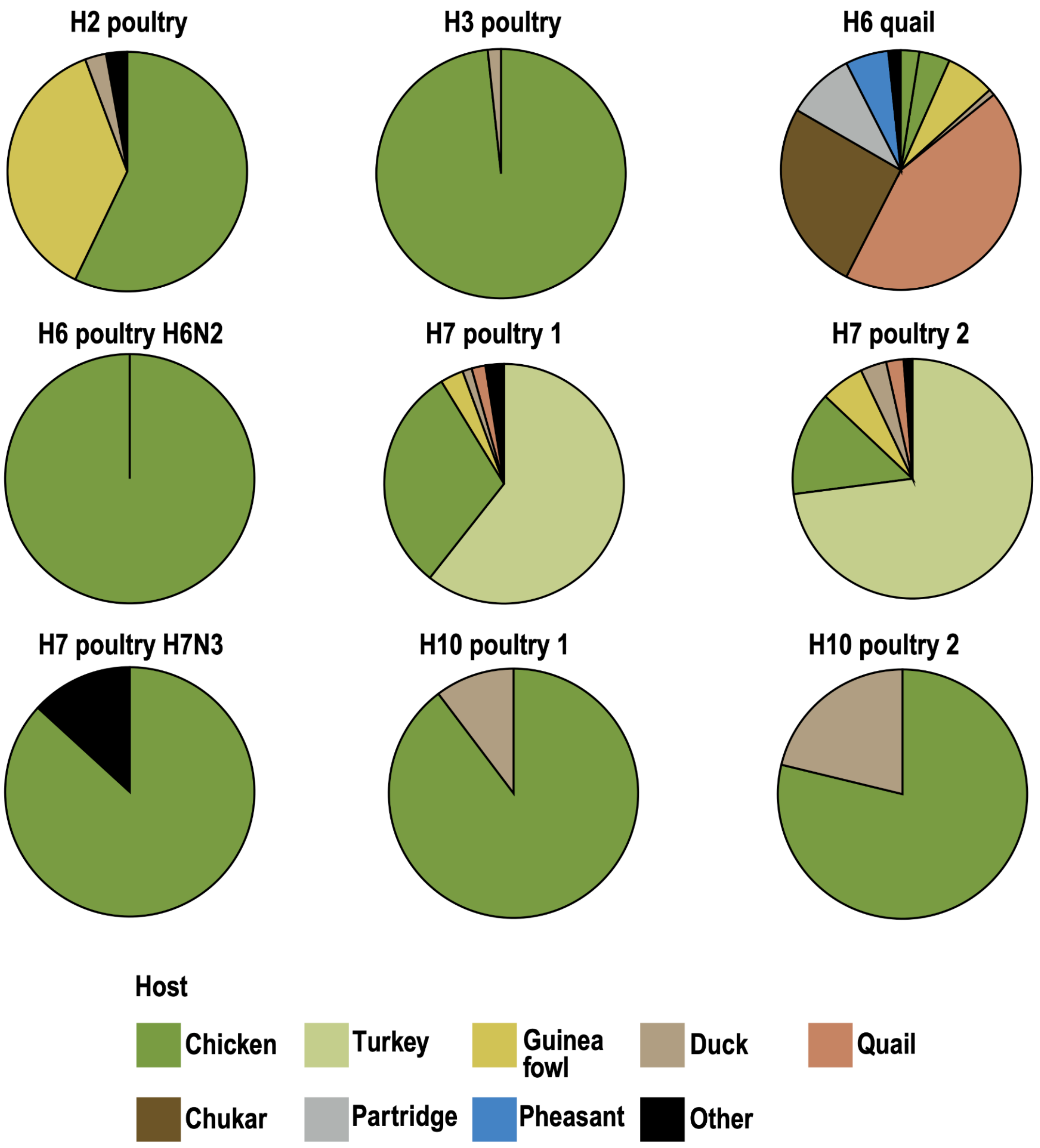


**Figure S12. Poultry IAV lineages are sampled from diverse poultry hosts.** The proportion of IAV sequences in each of the nine minor poultry IAV lineages (***Figures 3D-E***) is shown. Several lineages are predominantly isolated from chickens, while others are predominantly isolated from non-chicken hosts. As all of these lineages exhibit respiratory-like spectra (***Figures 3D-E***), this suggests that IAV undergoes predominantly respiratory infections and transmission in all poultry species.

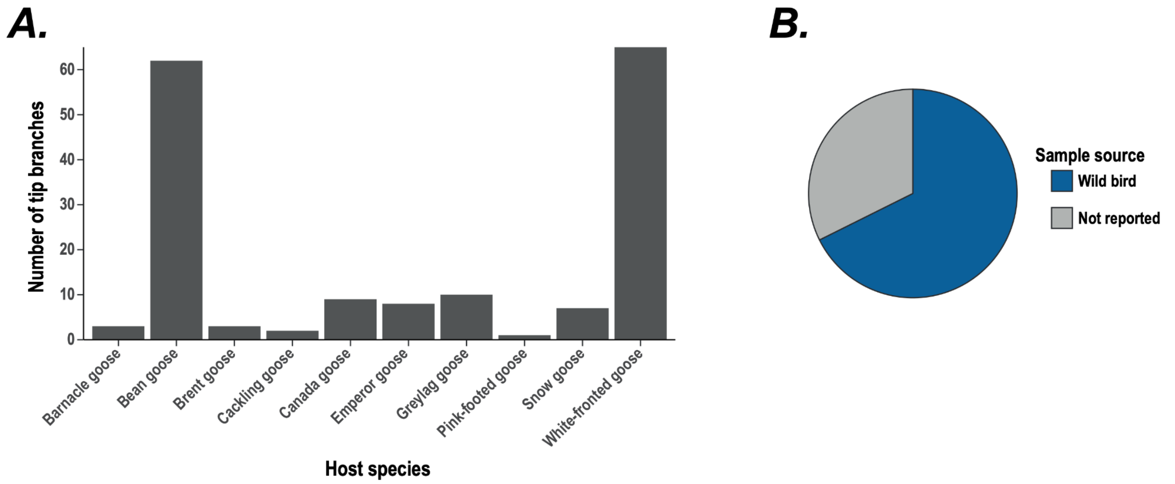


**Figure S13. The goose (*Anser*) tip branch spectra are calculated on viruses isolated from wild birds.** As major IAV lineages established in domestic Anseriformes exhibit a respiratory-like spectrum (***Figure 2C***), we checked that the respiratory tip phylogenetic branch spectrum in geese (***Figure 3F***) is not due to these birds being domestic. (**A**) The geese tip branches are predominantly isolated from bean goose and white-fronted goose in geographical areas where these species occur naturally and in publications sampling wild birds. (**B**) We employed original literature to review the status of geese from which IAV was isolated. All of the birds for which this was reported (68%) are wild birds. This demonstrates that the tip branch spectrum was reconstructed in wild birds and therefore that IAV is respiratory in wild geese.

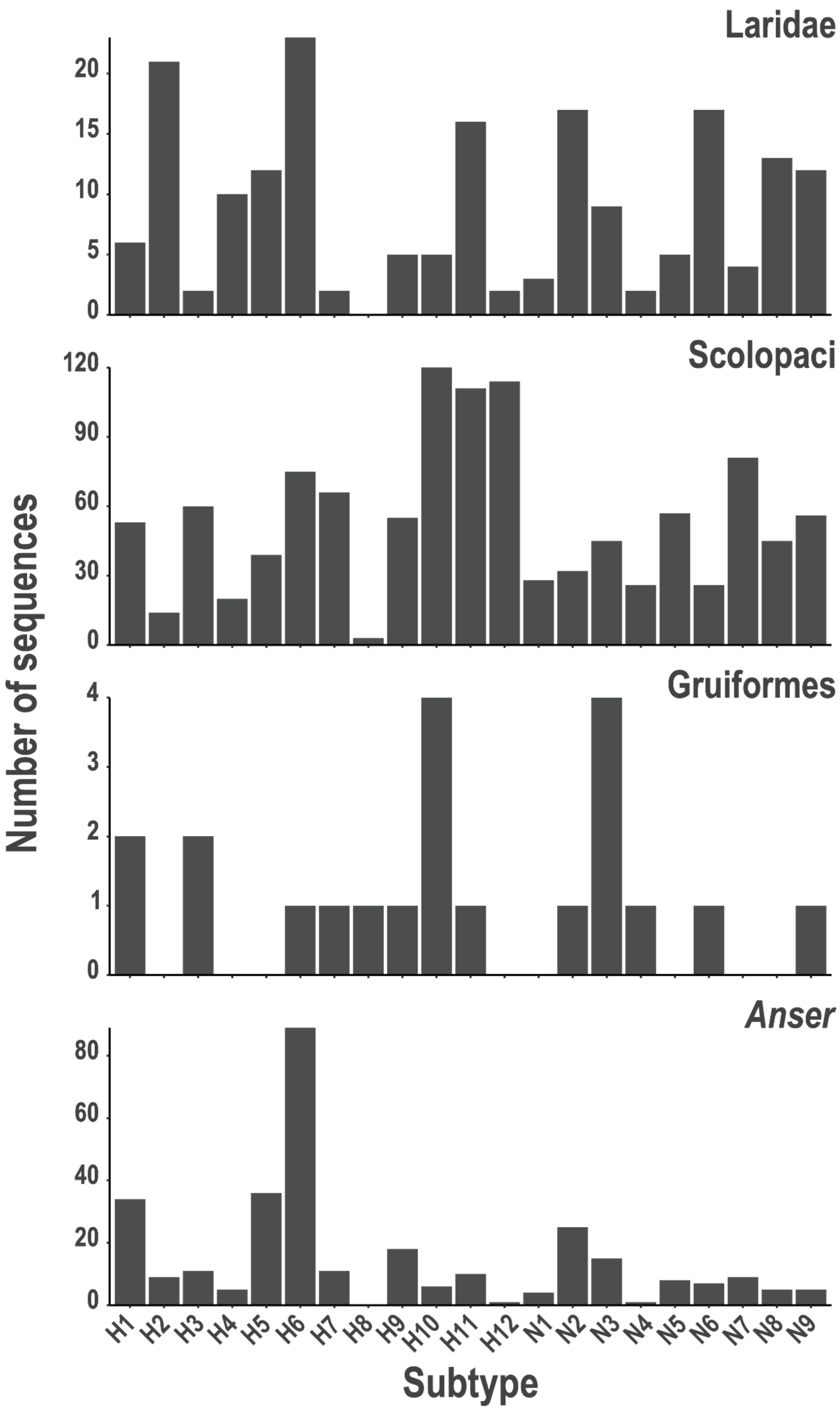


**Figure S14. IAV sequences are broadly distributed across subtypes in wild waterbirds with respiratory mutational spectra.** The number of IAV sequences in each subtype is shown for each of the major wild waterbird groups that exhibit respiratory-like mutational spectra.

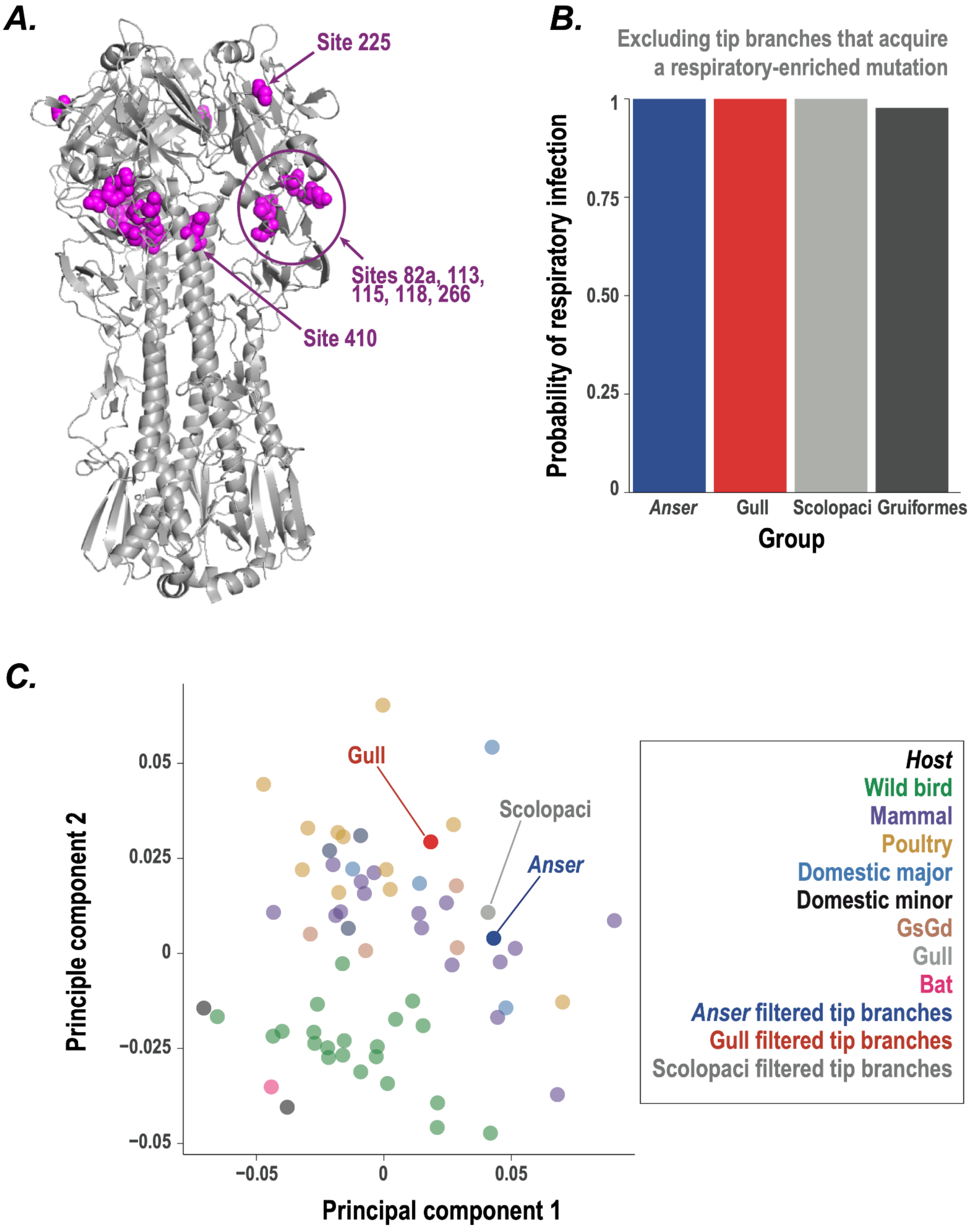


**Figure S15. IAV HA mutations assist with respiratory tropism but are not required for respiratory infection.** (**A**) IAV HA structure (PDB accession 4FNK) labelled with eight sites where mutations are enriched on tip branches leading to wild waterbird groups that exhibit respiratory mutational spectra. Site 532 is not shown as this site is not present in 4FNK. Sites 82a, 113, 115, 118 and 266 cluster together within the HA structure. (**B**-**C**) The probability of respiratory infection over gastrointestinal infection (**B**) and clustering (**C**) for tip branches sampled from wild waterbird taxonomic groups, excluding tip branches that acquire mutations at one or more of the sites associated with respiratory infection. Each group still exhibits strong support for respiratory transmission, showing that the respiratory-enriched mutations are not required for respiratory infection.

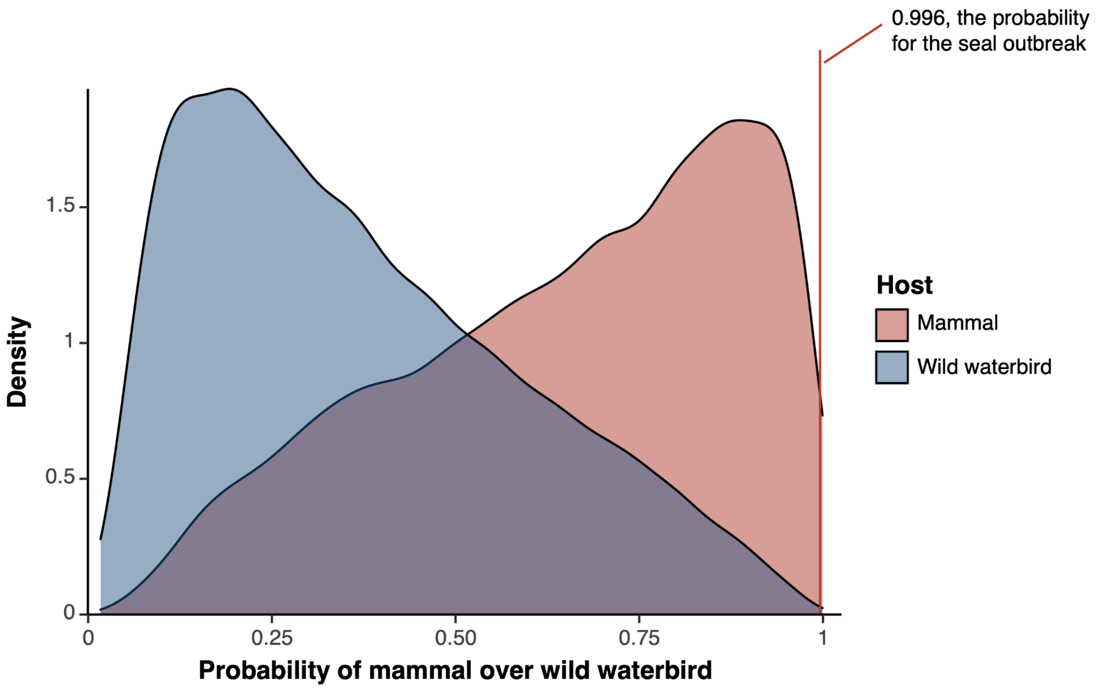


**Figure S16. Inference of transmission route is possible with 53 mutations.** To determine whether transmission route can be inferred with a small number of mutations, we randomly subsampled each wild waterbird and mammal spectrum within the 61 lineage dataset to 53 mutations (the number of mutations in the seal H10N7 outbreak lineage) 1000 times. We calculated the probability of each subsampled spectrum being generated by a mammalian spectrum over a wild waterbird spectrum using the likelihood-based classifier and here show the distribution of those probabilities separated by whether the lineage spectrum is from mammals or wild waterbirds. The probability of mammal over wild waterbird for the seal H10N7 outbreak is indicated.

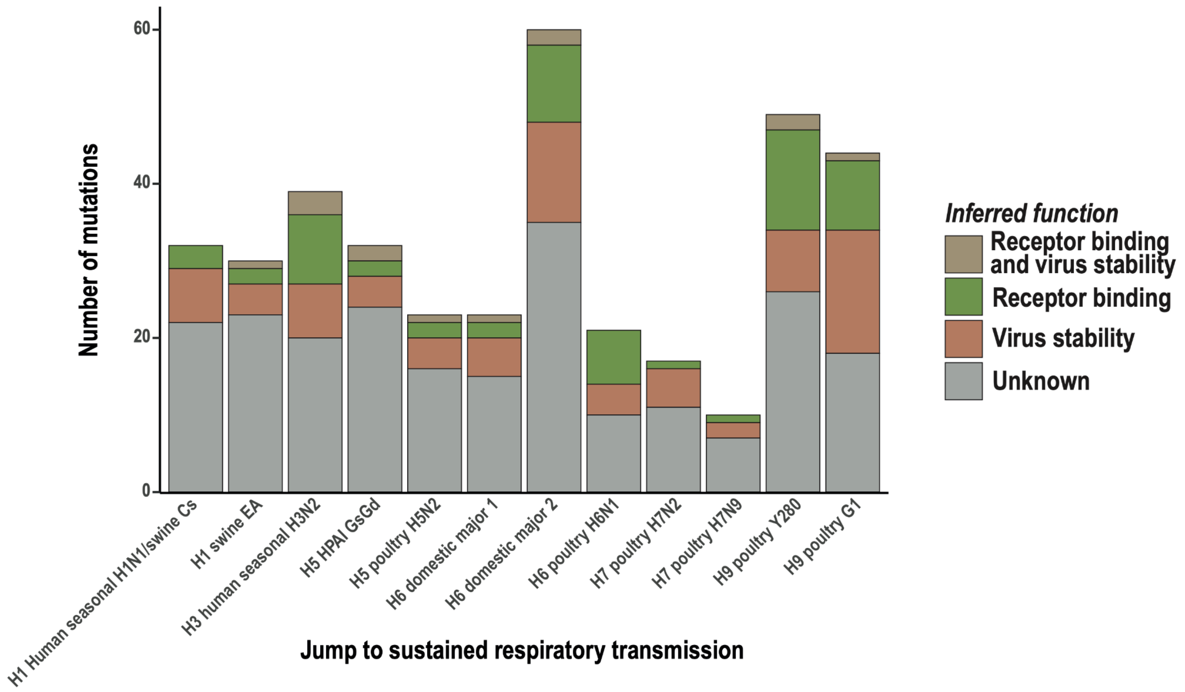


**Figure S17. Mutations inferred to impact receptor binding and stability are acquired upon each change to sustained respiratory transmission.** We inferred mutations associated with mutational spectrum shifts from gastrointestinal to respiratory (***Figure 4A***) and inferred the likely function of each mutation using DMS data (see ***Methods***). The number of mutations with each inferred function is shown for each shift to respiratory mutational spectrum. Each spectrum shift is associated with mutations inferred to impact receptor binding and mutations inferred to impact virus stability.

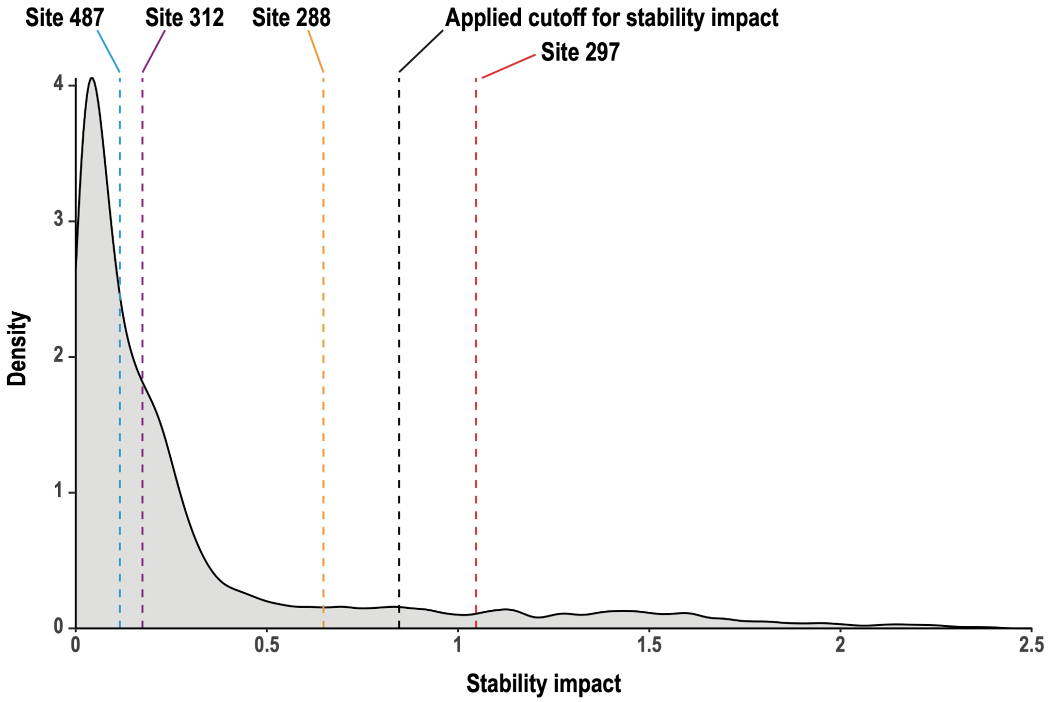

**Figure S18. Stability impacts of sites that mutate convergently leading to sustained respiratory transmission across multiple subtypes.** The distribution shows the magnitude of impact on stability of all mutations tested within DMS data in an H5 HPAI GsGd background (*59*). The maximum single mutation stability impact of the four non-RBD sites we identify as mutating convergently across subtypes (***Figure 4B***) is shown. Site 297 is above the threshold we set for stability impact based on known stability mutations (see ***Methods***) while site 288 is close to the threshold, supporting that mutations within this region of HA can impact virus stability.

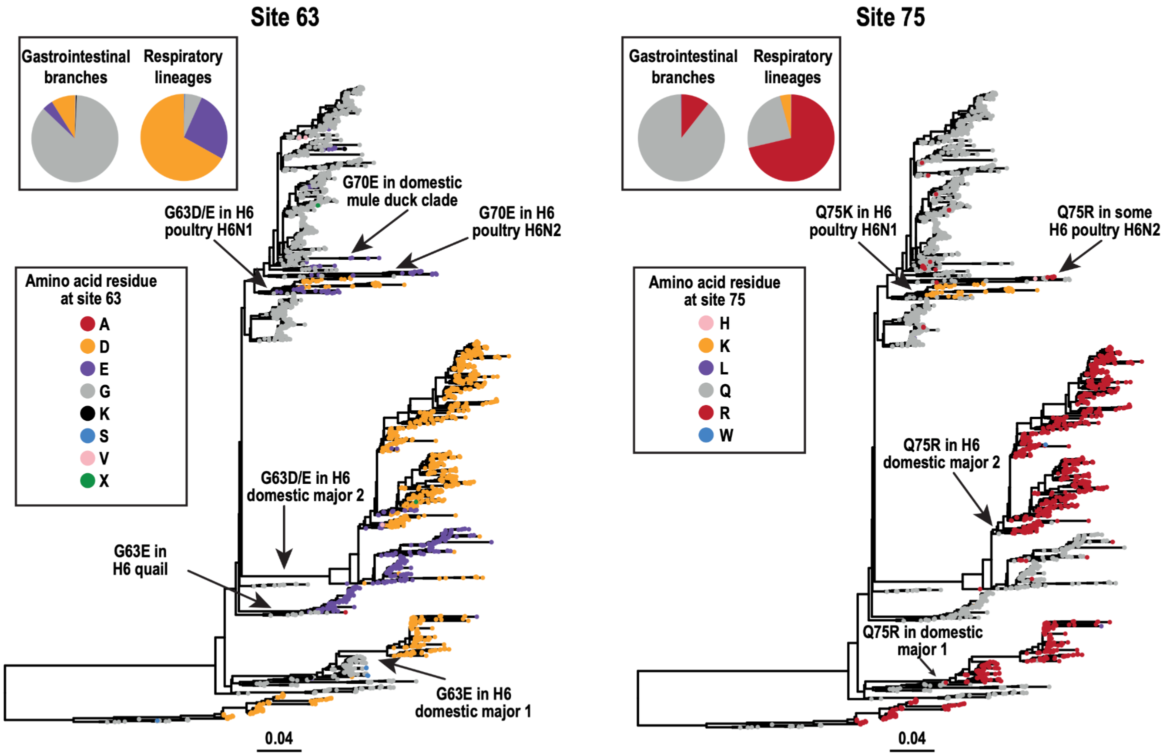


**Figure S19. Mutations at sites 63 and 75 are associated with mutational spectrum change in the H6 subtype.** The H6 phylogenetic tree is coloured by the amino acid residue at site 63 (left panel) or site 75 (right panel). Mutations at each site leading to sustained respiratory lineages are labelled. The pie chart insets show the distribution of amino acid residues at the respective site in the wild waterbird gastrointestinal sequences and in lineages with spectrum evidence of sustained respiratory transmission.
