## Supplementary material for "Mutational spectra reveal influenza virus transmission routes and adaptation": Table S7

### SUPPLEMENTAL TABLE

#### Data Availability

| | |
|---|---|
| GISAID Identifier: | EPI_SET_251120bq |
| DOI: | https://doi.org/10.55876/gis8.251120bq |

All genome sequences and associated metadata in this dataset are published in GISAID's EpiFlu database. To view the contributors of each individual sequence with details such as accession number, Virus name, Collection date, Originating Lab and Submitting Lab and the list of Authors, visit EPI_SET_251120bq

#### Data Snapshot

EPI_SET_251120bq is composed of 76777 individual viruses.
The collection dates range from 1905-05-11 to 2023-10-14;
Data were collected in 172 countries and territories.
